# Restoring vascular endothelial autophagic flux reduces atherosclerotic lesions

**DOI:** 10.1101/2021.05.06.442901

**Authors:** Shruti Chatterjee, Marouane Kheloufi, Stephane Mazlan, Xavier Loyer, Timothy A. McKinsey, Chantal M. Boulanger, Pierre-Michaël Coly

## Abstract

Atherosclerotic lesions preferentially develop in arterial areas exposed to low shear stress, where endothelial cells express a pro-inflammatory, apoptotic, and senescent phenotype. Endothelial cells exposed to atheroprone low shear stress present a defective autophagic flux, which favors a pro-inflammatory phenotype and the formation of atherosclerotic lesions. We tested the hypothesis that HDAC6 inhibition could restore adequate levels of autophagy in endothelial cells exposed to low shear stress. We found that blocking HDAC6 activity, either by pharmacological inhibition (Tubastatin-A) or genetic approaches (shHDAC6), raised levels of acetylated α-tubulin, as well as LC3-II/I ratio, LC3 puncta area, nuclear TFEB translocation and autophagic flux in cultured endothelial cells exposed to low shear stress. This effect was associated with a reduced expression of inflammatory markers (ICAM-1, VCAM-1 and MCP-1) in TNF-α-stimulated cells. Impaired endothelial autophagic flux was restored in the aortic arch (atheroprone conditions) of *HDAC6*^*-/-*^mice, when compared to control littermates. Atherosclerotic plaque size was significantly decreased in the aortic arch of chimeric *HDAC6*^*-/-*^*/ApoE*^*-/-*^ mice, transplanted with *HDAC6*^*+/+*^*/ApoE*^*-/-*^ bone marrow, when compared to *HDAC6*^*+/+*^*/ApoE*^*-/-*^ littermate controls. Taken together, these results indicate that HDAC6-inhibition may be an interesting strategy to restore endothelial autophagic flux, to help promote an atheroprotective endothelial phenotype despite unfavorable shear stress conditions.

## Introduction

Over the years, several studies have shown that atherosclerotic plaques form in predisposed areas such as arterial bifurcations, or the inner curvature of aortic bends. In these regions, endothelial cells are exposed to disturbed blood flow, which exerts low levels of shear stress (SS) on the vessel wall [1]–[3]. Conversely, cells exposed to laminar flow, which generates high SS, exhibit an anti-inflammatory, anti-senescent and anti-apoptotic phenotype, resulting in reduced plaque formation. Low SS triggers the expression of various inflammatory markers, such as adhesion proteins and chemokines [4], [5]. This in turn favors the recruitment of leukocytes to the lesion site and contributes to the initial stages of plaque formation. While the mechanisms by which SS regulates the endothelial phenotype have yet to be fully elucidated, our group has previously demonstrated that a modulation of endothelial macroautophagy (hereafter referred to as autophagy) is a determining factor [6].

Autophagy is an evolutionarily conserved, cytoprotective mechanism, which helps degrade and recycle damaged organelles and proteins, to maintain cellular homeostasis [7]. As such, autophagy occurs at a basal rate under physiological conditions, and is amplified under stress conditions, such as nutrient starvation, to promote cell survival [8]. Defects in the autophagic process have been reported in multiple pathologies, including cancer, neurodegenerative, auto-immune and cardiovascular diseases [9]–[12]. Often, the defect lies in the crucial autolysosome formation step, wherein autophagosomes fuse with lysosomes to allow the degradation of their content [11], [13]. Low SS has been shown to inhibit autophagic flux in endothelial cells, which then contributes to the onset of a pro-inflammatory and senescent phenotype [6], [14], [15]. It is unknown at the moment whether or not increasing endothelial autophagic flux in atheroprone areas of the vasculature would prevent endothelial inflammation and subsequent atherosclerotic plaque development in these unfavorable shear stress conditions.

While numerous factors can regulate autophagy, studies have highlighted the importance of several post-translational tubulin modifications and their effects on microtubule dynamics and cellular trafficking [16]. In addition to concentrating signaling pathways that stimulate autophagosome formation, microtubules are integral to the movement of pre-autophagosomal structures, as well as the transport of mature autophagosomes towards lysosomes [16]–[18]. In that sense, microtubules stabilized through α-tubulin acetylation favor the recruitment and walking of motor proteins, resulting in enhanced cellular trafficking [18]. Endothelial cells exposed to high SS are characterized by a profound stabilization of microtubules, associated with increased acetylated α-tubulin when compared to cells cultured under static conditions [19]. This may, in part, explain why endothelial cells exposed to high SS have an increased autophagic activity. Interestingly, the level of endothelial α-tubulin acetylation and microtubule stabilization are regulated by cytosolic protein deacetylases and in particular by Histone Deacetylase 6 (HDAC6) [20], [21], whose phosphorylated form remains active in the cytosol [22], [23]. Other pathways beside microtubule stabilization also regulate endothelial autophagic flux; previous studies have shown that endothelial atheroprotective shear stress activates transcription factor EB (TFEB), leading to increased lysosomal biogenesis and autophagic flux, and subsequent anti-inflammatory endothelial phenotype [24], [25]. TFEB activity and its nuclear localization are regulated by posttranscriptional modifications such as acetylation by HDAC6 [26] and HDAC inhibition in experimental kidney disease leads to TFEB activation and prevents the accumulation of misfolded aggregates [27].

Based on these data, we hypothesized that HDAC6 inhibition could restore adequate levels of autophagy in endothelial cells exposed to atheroprone low SS. In this study, we found that HDAC6 activity was more potent in atheroprone than atheroprotected endothelial cells. In addition, in vitro HDAC6 targeting using a selective inhibitor, Tubastatin-A, or an shRNA, increased autophagic flux and reduced expression of inflammatory markers ICAM-1, VCAM-1 and MCP-1 in an autophagy-dependent manner. Subsequent *in vivo* experiments demonstrated that knocking out *HDAC6* in hypercholesteremic mice reduces the formation of atherosclerotic lesions in otherwise predisposed areas in the aorta.

## RESULTS

### HDAC6 activity is increased in endothelial cells exposed to low SS

HDAC6 deacetylase activity was determined as total deacetylase activity sensitive to a known HDAC6 inhibitor, Tubastatin-A (3 µM) [28]; this activity was significantly higher in endothelial cells exposed to LSS than in cells exposed to HSS for 24 hours (Figure 1A). The expression and localization of both the canonical and phosphorylated forms of HDAC6 were analyzed by immunofluorescence in HUVECs exposed to different shear stress levels for 24 hours (Figure. 1B). While no difference in the staining intensity of total-HDAC6 was observed, HUVECs exposed to low SS exhibited higher levels of phosphorylated-HDAC6, the active HDAC6 isoform, compared to cells exposed to high SS conditions (Figure.1C and 1D). Taken all together, these results indicate that endothelial cells exposed to atheroprone low SS have a higher HDAC6 activity than cells exposed to atheroprotective high SS.

**Figure 1:**
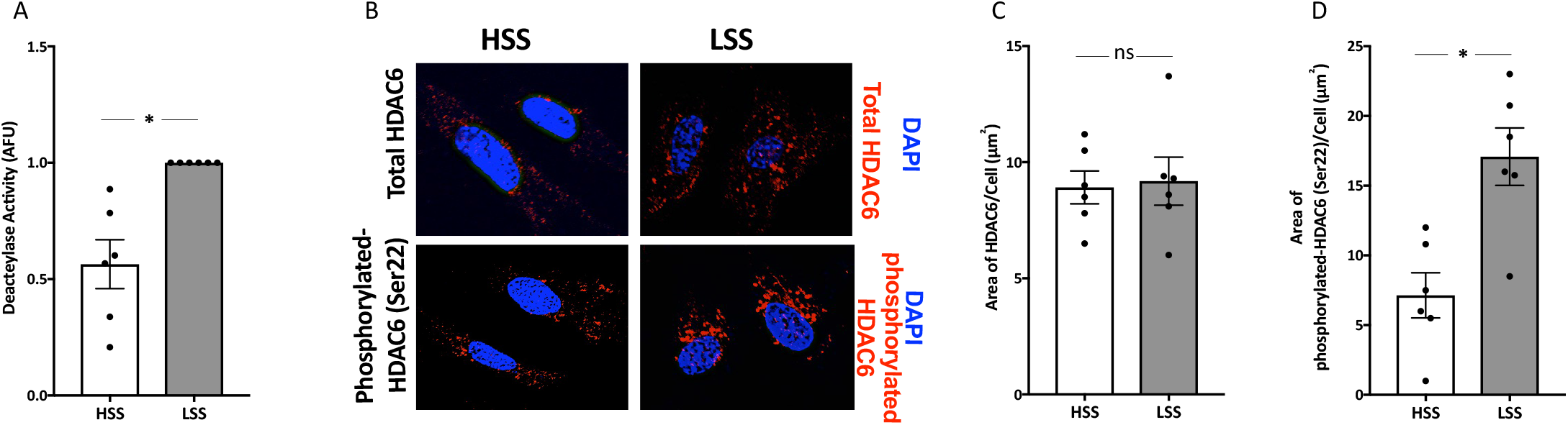
Regulation of HDAC6 activity by shear stress. **(A**) Analysis of HDAC6 activity in HUVECs exposed to high SS or low SS for 24 hours. Data represent means ± SEM of 6 independent experiments, normalized to low SS conditions. \**P* ≤ *0*.*05* (Wilcoxon test). **(B)** Representative images and analysis of expression of HDAC6 and phosphorylated-HDAC6 by immunofluorescence. HUVECs exposed to high SS (HSS; 20 dyn/cm^2^) or low SS (LSS; 2 dyn/cm^2^) for 24 hours, were labeled with HDAC6 (red), or phosphorylated-HDAC6 antibody (red), and the area/cell (µm^2^) was analyzed (blue=DAPI). Quantification of total HDAC6 **(C)** and phosphorylated HDAC6 **(D)** area in endothelial cells exposed to either high SS or low SS. Data represent means ± SEM of 6 independent experiments, normalized to low SS conditions. ns, not statistically different; \**P* ≤ 0.05 (Wilcoxon test).

As expected from McCue et al. [29], acetylated α-tubulin levels were higher in endothelial cells exposed to high SS, when compared to low SS (Supplementary Figure 1). Tubastatin-A markedly increased levels of acetylated α-tubulin in cells exposed to low SS (Supplementary Figure. 2A and 2B). Additionally, acetylated α-tubulin displayed a more organized pattern in Tubastatin-A-treated cells (Suppl. Figure. 2C and 2D). These data indicate that inhibition of HDAC6 leads to stabilization of a polymerized α-tubulin network in endothelial cells exposed to low SS.

### HDAC6 inhibition restores autophagic activity in endothelial cells exposed to low SS

We next evaluated whether HDAC6 inhibition would stimulate autophagic activity in endothelial cells exposed to low SS. We first assessed autophagic activity by measuring LC3-II/I ratio and observed that, in accordance with our previous findings, cells exposed to low SS for 24h displayed a lower LC3-II/I ratio than cells exposed high SS. Treatment with Tubastatin-A, in low SS conditions, increased the LC3-II/I ratio to approximately twice that of control conditions (Figure 2A and 2B). Similar results were obtained when expressing LC3 as a ratio of LC3-II to GAPDH. Interestingly, Tubastatin-A did not modify levels of other essential autophagy proteins, such as ATG5 (Supplementary Figure. 3A and 3B) and p62 (Supplementary Figure. 3C and 3D). We obtained comparable data when measuring the area of LC3-positive puncta per cell by immunostaining. LC3 staining was consistently less abundant in endothelial cells exposed to low SS. However, treating cells with Tubastatin-A elevated the area of LC3-positive structures per cell to levels similar to high SS conditions (Figure. 2C and 2D).

**Figure 2:**
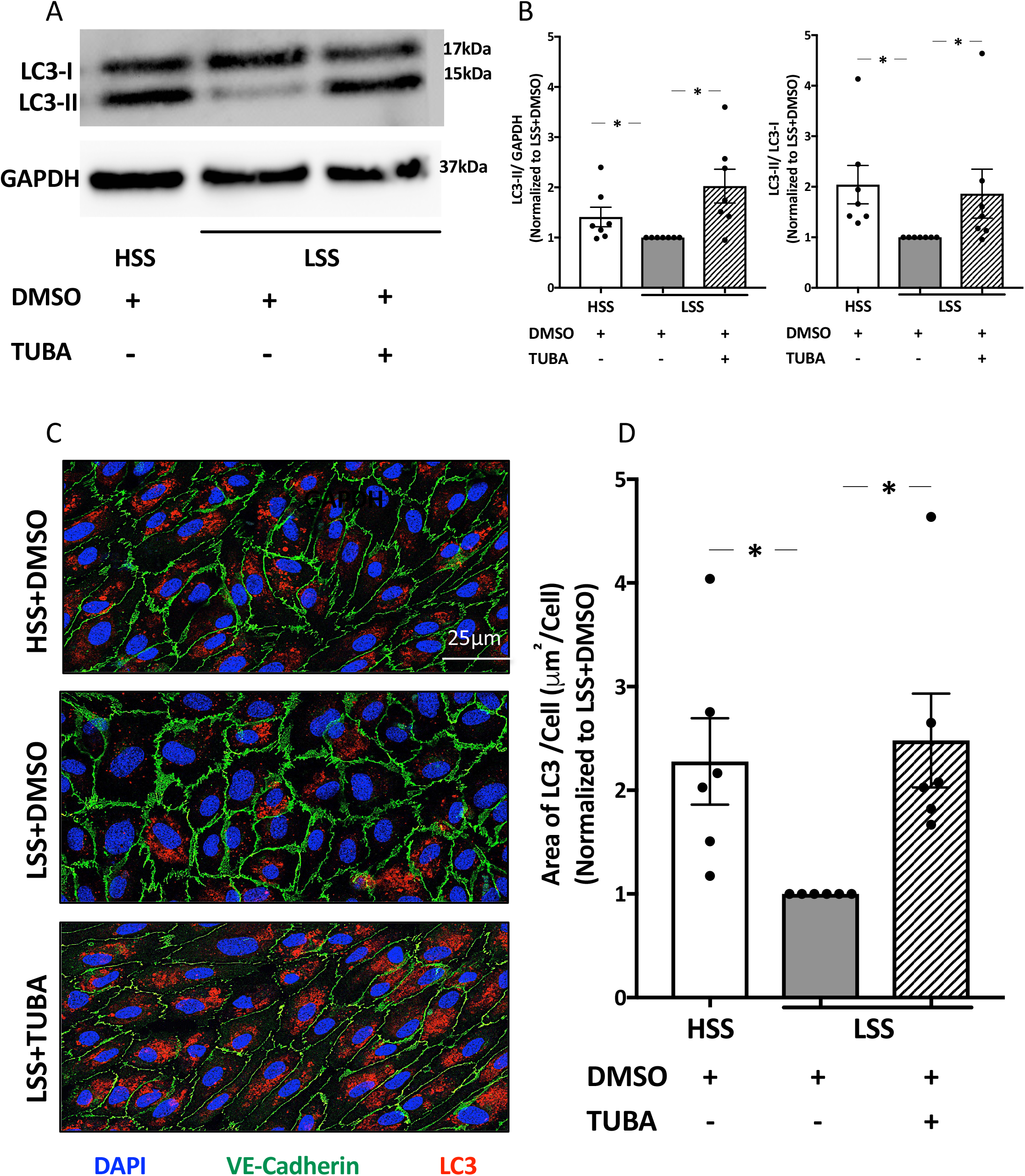
Tubastatin-A treatment increases endothelial autophagy. **(A)** Western blot analysis of LC3 in HUVECs exposed to either high SS (HSS; 20 dyn/cm^2^) or low SS (LSS; 2 dyn/cm^2^) and treated with either vehicle (DMSO: 0.1 µL/mL) or Tubastatin-A (TUBA; 3 µM) for 24 hours. **(B)** Quantification of the LC3-II/I (left) and LC3-II/GAPDH (right) ratios; data represent means ± SEM of 6 independent experiments normalized to low SS + DMSO. **(C)** Representative confocal microscopy images of HUVECs exposed to either high SS or low SS and treated with either vehicle (DMSO: 0.1 µL/mL) or Tubastatin-A (TUBA; 3 µM) for 24 hours (blue: DAPI, green: VE-cadherin, red: LC3). **(D)** Quantification of LC3 area per cell; data represent means ± SEM of 6 independent experiments normalized to low SS + DMSO. \**P* ≤ *0*.*05* (Wilcoxon test).

To confirm that Tubastatin-A stimulates endothelial autophagy, we used a tandem LC3-B fluorescently tagged with mRFP and GFP. In this assay, autophagosomes emit a yellow signal (mRFP-GFP-LC3), and their maturation into autolysosomes is attested by a red signal due to quenching of the GFP fluorescence in low pH environments (Figure 3A). We therefore evaluated autophagic flux by quantifying the ratio of autolysosomes/autophagosomes per cell (Figure 3B). As previously demonstrated, endothelial cells exposed to high SS had approximately twice the number of autolysosomes than those exposed to low SS conditions, while the number of autophagosomes was not different between the two experimental conditions (Figure 3C and 3D). This indicated that maturation of autophagosomes into autolysosomes was impaired under low SS conditions. Treatment with Rapamycin (1 µM), a known autophagy inducer, resulted in a slight decrease in the number of autophagosomes, and an increase in the number of autolysosomes. A similar trend was observed in cells exposed to Tubastatin-A under low SS conditions, suggesting that HDAC6 inhibition stimulates autophagic flux in endothelial cells exposed to low SS. However, these results could also be explained by an increase in autophagosome biogenesis. To test this hypothesis, we examined the effects of Tubastatin-A in combination with chloroquine, a lysosomotropic agent that neutralizes the acidic pH of lysosomes, thereby blocking the fusion between autophagosomes and lysosomes [30]. As expected, treating cells with chloroquine (300μM) resulted in an increase in the number of autophagosomes (Figure 3E and 3F). In the presence of chloroquine, Tubastatin-A failed to elevate the number of autolysosomes, but increased the number of autophagosomes, which suggests an acceleration of autophagosome formation (Figure 3G).

**Figure 3:**
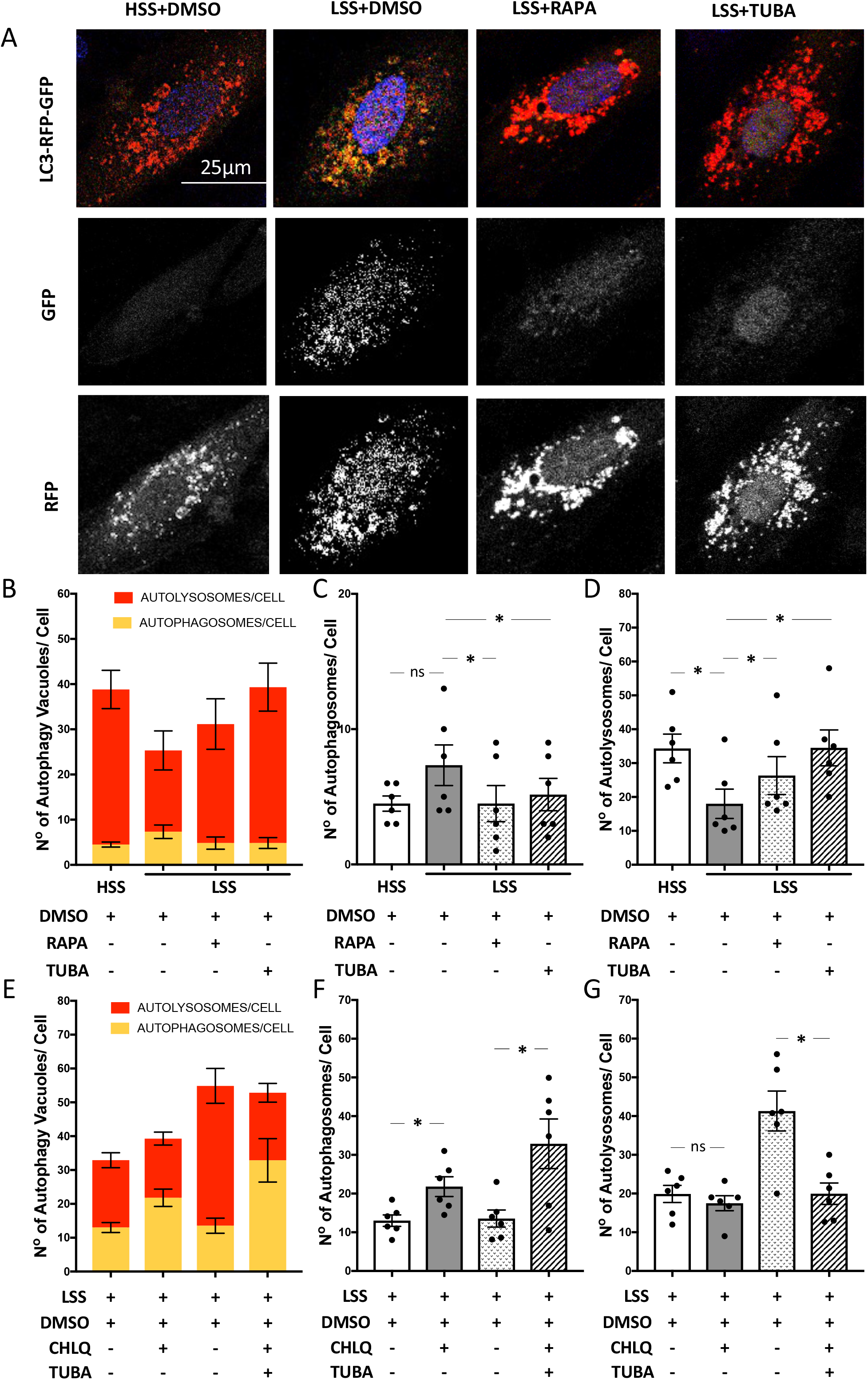
Autophagic flux is restored in HUVECs exposed to low SS and treated with Tubastatin-A. **(A)** Representative confocal microscopy images of HUVECs infected with lentiviruses carrying the tandem mRFP-GFP-LC3 plasmid and exposed to high SS (HSS; 20 dyn/cm^2^) or low SS (LSS; 2 dyn/cm^2^) for 24 hours, in the presence or absence of Tubastatin-A (TUBA; 3 µM) or Rapamycin (RAPA; 1 µM). Autophagosomes and autolysosomes are denoted by yellow and red signals, respectively. Scale bar: 25 µm. **(B)** Quantification of the number of autophagy vacuoles per cell. Yellow bars correspond to the number of autophagosomes per cell and red bars correspond to the number of autolysosomes per cell. HUVECs were exposed to either high SS or low SS and treated with either vehicle (DMSO at 0.1 µL/mL), Rapamycin (RAPA; 1µM) or Tubastatin-A (TUBA; 3 µM) for 24 hours. Quantification of the number of autophagosomes **(C)** and autolysosomes **(D)** per cell. **(E)** Quantification of the number of autophagy vacuoles per cell in HUVECs exposed to low SS and treated with Chloroquine (CHLQ; 300µM) and either vehicle (DMSO at 0.1 µL/mL) or Tubastatin-A (TUBA; 3 µM) for 24 hours. Quantification of the number of autophagosomes **(F)** and autolysosomes **(G)** per cell in HUVECs exposed to low SS and treated with Chloroquine (CHLQ; 300µM) and either vehicle (DMSO at 0.1 µL/mL) or Tubastatin-A (TUBA; 3 µM) for 24 hours). Data represent means ± SEM of 6 independent experiments in which over 100 cells were analyzed per condition. ns, not statistically different; \**P* ≤ 0.05 (Wilcoxon test).

As lysosomal biogenesis is an important factor in promoting efficient autophagic flux, we assessed the effect of Tubastatin-A on Transcriptional Factor EB (TFEB) activity, a well-known regulator of lysosomal biogenesis [31], [32]. Tubastatin-A not only upregulates the total expression of lysosomal transcription factor, TFEB (Supplementary Figure 4A-C), but also counter-acts its sequestration in the cytosol (phosphorylated TFEB) [33] by promoting its translocation to the nucleus (Supplementary Figure 4E-H).

We then examined lysosomal biogenesis by evaluating the expression of lysosomal marker, LAMP-1, in these experimental conditions where Tubastatin-A increased TFEB activity. We observed no significant increase in LAMP-1 expression under Tubastatin-A treatment (Supplementary Figure. 4A and Supplementary Figure. 4D) [34], indicating that under the present experimental conditions, Tubastatin-A favors autophagic flux in autophagy-deficient conditions, without facilitating the formation of lysosomes via increasing TFEB activity.

### HDAC6 inhibition reduces oxidative stress and endothelial inflammation caused by exposure to low SS conditions

Our group has previously demonstrated that the defect in endothelial autophagy, which occurs in low SS areas, causes inflammation and favors the development of atherosclerotic lesions [6]. Since Tubastatin-A treatment restored autophagic flux in endothelial cells exposed to low SS, we wondered if this would also hamper the pro-inflammatory consequences of these atheroprone conditions. To induce inflammation, cells were treated with TNF-α, at 1ng/mL, for the last 12 hours of the shear stress experiment, resulting in a pronounced elevation of ICAM-1 (Figure 4A and 4C) and VCAM-1 (Figure 4B and 4D) expression in cells exposed to low SS expressed, as compared to cells exposed to high SS. This response was significantly reduced in the presence of Tubastatin-A. We next measured the concentration of MCP-1 found the HUVEC supernatant, by ELISA. Endothelial cells exposed to low SS released approximately 5 times more MCP-1 than cells exposed to high SS conditions. Here also, adding Tubastatin-A to cells, markedly reduced levels of MCP-1 released by cells exposed to low SS (Figure 4E). Interestingly, Tubastatin-A also suppressed the TNF-α-induced elevation in phospho-NF-κB levels (Supplementary Figure 5A and Supplementary Figure 5B) and in reactive oxygen species levels (Supplementary Figure 5C and Supplementary Figure 5D).

**Figure 4:**
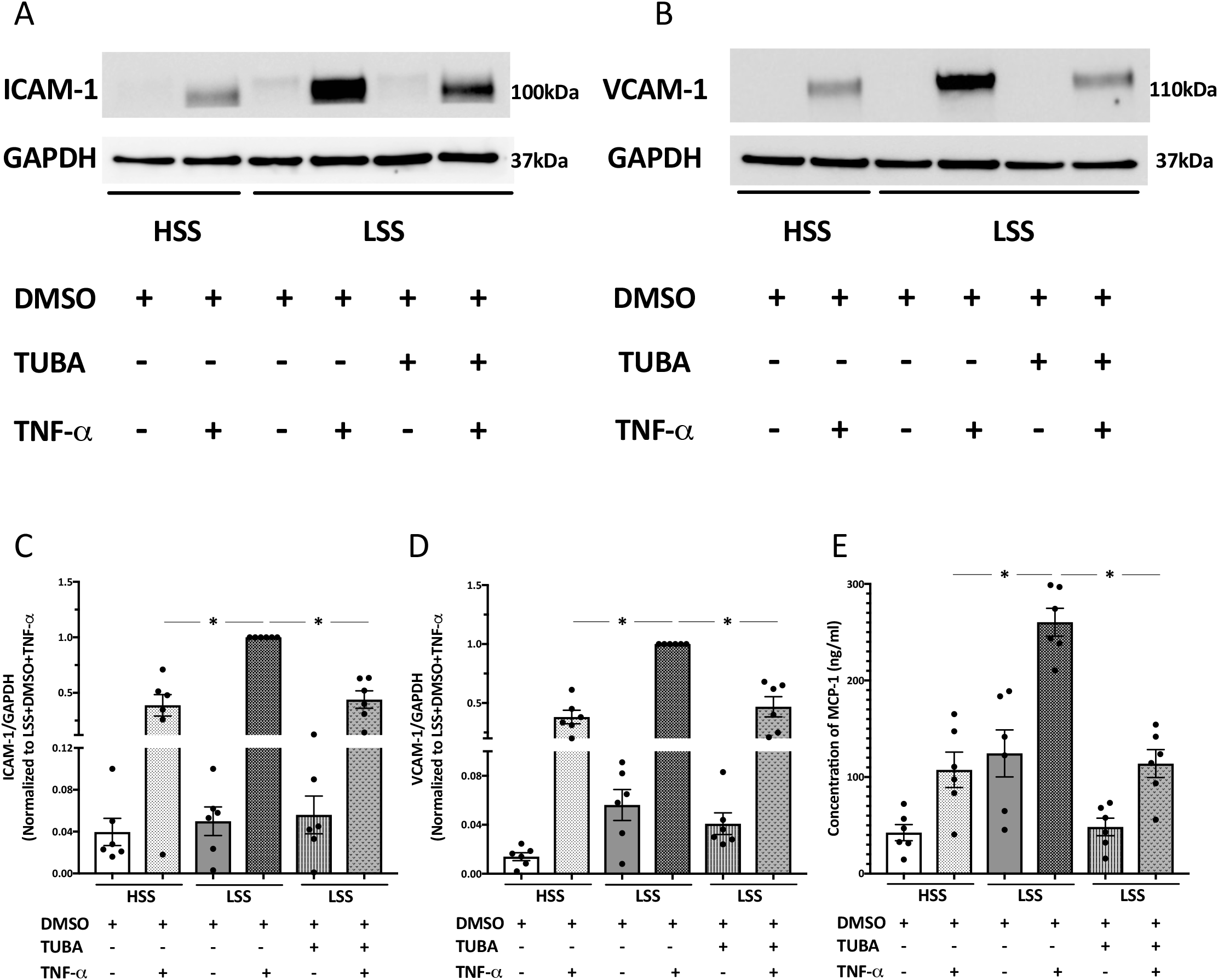
Tubastatin-A has anti-inflammatory effect on endothelial cells exposed to low shear stress. **(A)** Western blot analysis of ICAM-1 in HUVECs exposed to high SS (HSS; 20 dyn/cm^2^) or low SS (LSS; 2 dyn/cm^2^) and treated with either vehicle (DMSO at 0.1 µL/mL) or Tubastatin-A (TUBA; 3 µM) for 24 hours, in the presence or absence of TNF-α (1 ng/mL). **(B)** Western blot analysis of VCAM-1 in HUVECs exposed to high SS (HSS; 20 dyn/cm^2^) or low SS (LSS; 2 dyn/cm^2^) and treated with either vehicle (DMSO at 0.1 µL/mL) or Tubastatin-A (TUBA; 3 µM) for 24 hours, in the presence or absence of TNF-α (1 ng/mL). **(C)** Quantification of the ICAM-1/GAPDH ratio; data represent means ± SEM of 6 independent experiments normalized to low SS + DMSO + TNF-α. **(D)** Quantification of the VCAM-1/GAPDH ratio; data represent means ± SEM of 6 independent experiments normalized to low SS + DMSO + TNF-α. **(E)** ELISA analysis of MCP-1 levels released in the conditioned media of HUVECs exposed to high SS (HSS; 20 dyn/cm^2^) or low SS (LSS; 2 dyn/cm^2^) and treated with either vehicle (DMSO at 0.1 µL/mL) or Tubastatin-A (TUBA; 3 µM) for 24 hours, in the presence or absence of TNF-α (1 ng/mL). Data represent means ± SEM of 6 independent experiments. \**P* ≤ 0.05 (Wilcoxon test).

To demonstrate that Tubastatin-A effects were mediated by HDAC6 inhibition and not by unknown off-target mechanisms, we knocked down HDAC6 in HUVECs using a shRNA-carrying lentivirus. With this strategy, we observed a 30% reduction in the expression of HDAC6 (Figure 5A and 5B) associated with a 50% increase in acetylated α-tubulin levels (Figure 5C). HDAC6-knockdown also led to an increase in the LC3-II/I ratio (Figure 5D), while reducing ICAM-1 (Figure 5E) and VCAM-1 (Figure 5F) expression as well as MCP-1 release (Figure 5G) in TNFα-treated cells. Taken together, these data show that inhibiting HDAC6 activity restores endothelial autophagy, and reduces reactive oxygen species accumulation, NF-κB activation and subsequent endothelial inflammation in cells exposed to low SS.

**Figure 5:**
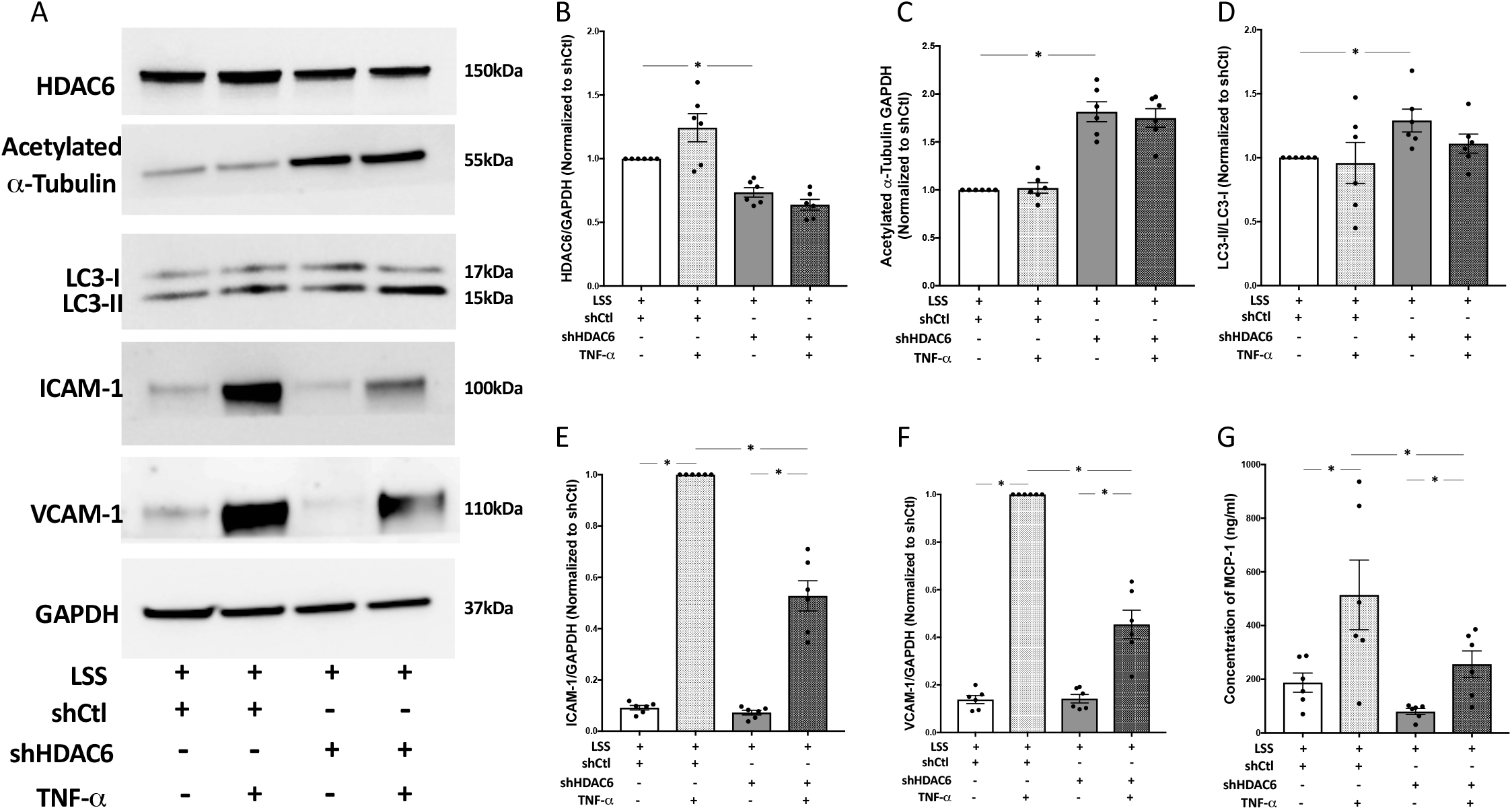
Knockdown of HDAC6 increases endothelial autophagy and reduces inflammation. **(A)** Western blots analysis of HDAC6, acetylated α-tubulin, LC3, ICAM-1 and VCAM-1 expression in HUVECs transduced with either shControl (shCtl) or shHDAC6 lentiviruses and exposed to low SS (LSS; 2 dyn/cm^2^), in the presence or absence of TNF-α (1 ng/mL). Quantification of the HDAC6/GAPDH **(B)**, acetylated α-tubulin/GAPDH **(C)**, LC3-II/I **(D)**, ICAM-1/GAPDH **(E)** and VCAM-1/GAPDH **(F)** ratios; data represent means ± SEM of 6 independent experiments normalized to shCtl. **(G)** ELISA analysis of MCP-1 levels released in the conditioned media of HUVECs transduced with either shControl (shCtl) or shHDAC6 lentiviruses and exposed to low SS (LSS; 2 dyn/cm^2^), in the presence or absence of TNF-α (1 ng/mL). Data represent means ± SEM of 6 independent experiments. \**P* ≤ 0.05 (Wilcoxon test).

### The anti-inflammatory effects of HDAC6 inhibition are relayed by autophagy

To verify whether the anti-inflammatory effects of HDAC6-inhibition were linked to a restored autophagic flux, we assessed the effects of Tubastatin-A in cells deficient for autophagosome formation (i.e., HUVECs infected with a lentivirus carrying an ATG5 shRNA) resulting in lowered ATG5 protein levels (Figure. 6A) and LC3-II/I ratio (Figure 6B). Endothelial HDAC6 protein expression was significantly augmented in shATG5 conditions (Supplementary Figure 7A and 7B). Silencing ATG5 did not affect acetylated α-tubulin expression, either in the absence or the presence of Tubastatin-A (Supplementary Figure 7A and 7C). In ATG5-deficient endothelial cells, the anti-inflammatory effect of Tubastatin-A was significantly reduced, as indicated by the lower expression of ICAM-1 (Figure. 6C and Supplementary Figure 6A), VCAM-1 (Figure 6D and Suppl. Figure. 6B) and the impaired MCP-1 release (Figure 6E and Supplementary Figure 6C). These data show that the anti-inflammatory effects of HDAC6-inhibition depend, at least in part, on a functional autophagic activity.

**Figure 6:**
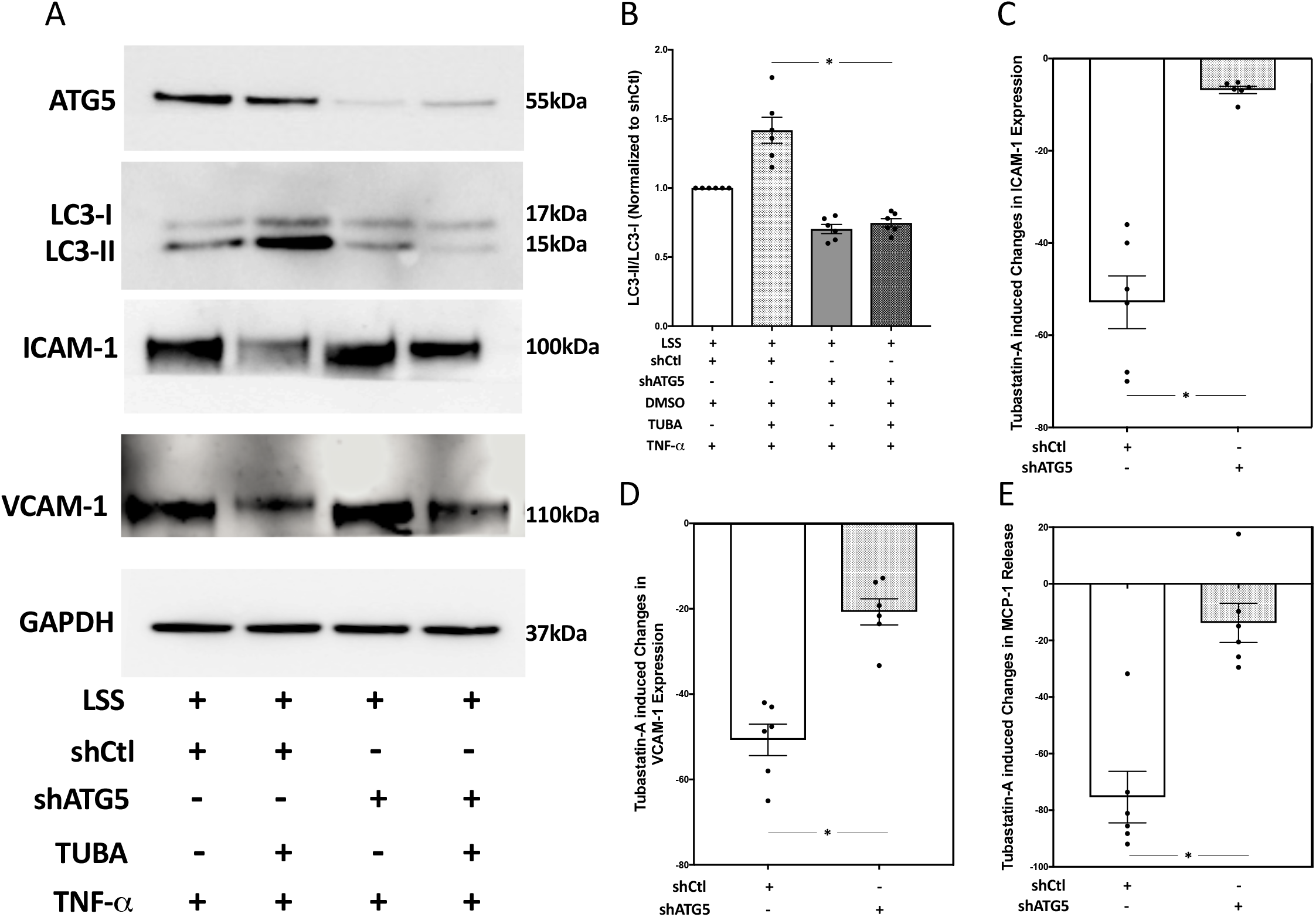
Deficient autophagy dampens the anti-inflammatory effect of Tubastatin-A. **(A)** Western blots analysis of ATG5, LC3, ICAM-1 and VCAM-1 expression in endothelial cells transduced with either shControl (shCtl) or shATG5 lentiviruses, exposed to low SS (LSS; 2 dyn/cm^2^), and treated with vehicle (DMSO at 0.1 μL/mL) or Tubastatin-A (TUBA; 3 μM) in the presence of TNF-α (1 ng/mL). **(B)** Quantification of the LC3-II/I ratio. Data represent means ± SEM of 6 independent experiments normalized to shCtl. Quantification of Tubastatin-A induced changes in the expression of ICAM-1 **(C)** and VCAM-1 **(D)**; and concentration of MCP-1 in the conditioned media of HUVECs **(E)**. Data represent means ± SEM of 6 independent experiments. ns, not statistically different; \**P* ≤ 0.05 (Wilcoxon test).

### Deletion of *HDAC6* enhances autophagic flux in murine aortic arch

To confirm the effect of *HDAC6* deletion on autophagic activity *in vivo*, we performed endothelial LC3 staining in *en face* murine aortas isolated from *HDAC6*^*-/-*^ */ ApoE*^*-/-*^ and littermate controls (*HDAC6*^*+/+*^*/ApoE*^*-/-*^). In control mice, we observed higher LC3 staining in the descending aorta (high SS), when compared to the aortic arch (low SS), corroborating previous results demonstrating that high SS activates endothelial autophagy [6] (Figure 7A and 7B). However, this difference no longer existed in endothelial cells from *HDAC6* ^*-/-*^*/ApoE* ^*-/-*^ mice. Interestingly, endothelial LC3 staining was significantly higher in the arch of *HDAC6*^*-/-*^ animals when compared to *HDAC6*^*+/+*^ controls (Figure 7A and 7B). Next, we evaluated endothelial autophagic flux by measuring the LC3 colocalization with the lysosomal marker LAMP2A. We observed that LC3-LAMP2A colocalization was significantly higher in the aortic arch of *HDAC6*^*-/-*^ animals compared to *HDAC6*^*+/+*^ control mice, which led us to infer that the absence of HDAC6 may favor promotion of autophagic flux in endothelial cells (Figure 7C and 7D). Taken all together, these results support the concept that deficiency in endothelial HDAC6 increases autophagic activity despite unfavorable shear stress conditions.

**Figure 7:**
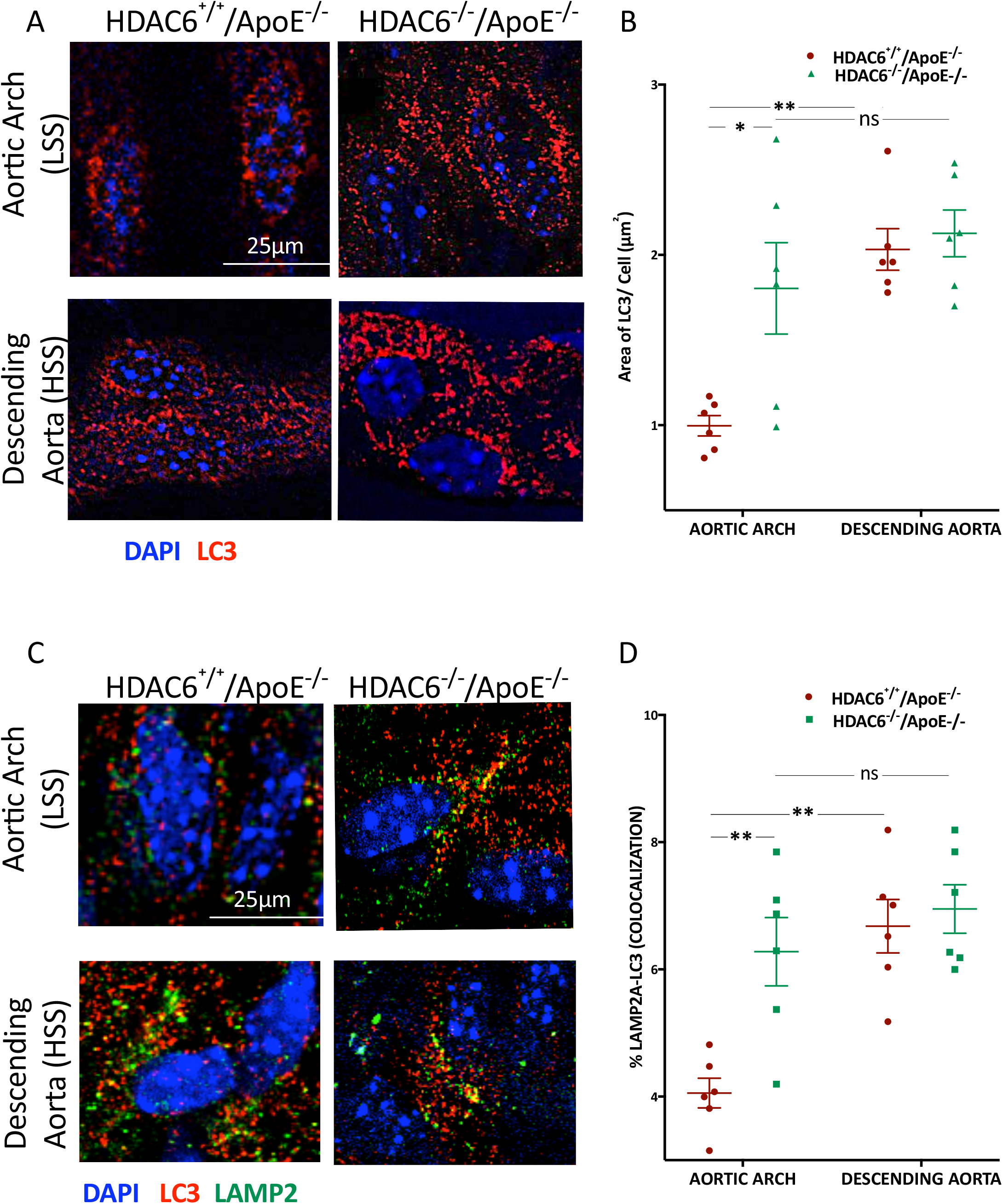
HDAC6 knockout increases autophagic flux in atheroprone areas of the aorta. **(A)** LC3 *en face* staining of the aorta of *HDAC6*^*-/-*^*/ApoE*^*-/-*^ mice and littermate controls *HDAC6*^*+/+*^*/ApoE*^*-/-*^; n=3 per group; (red, LC3; blue, DAPI). **(B)** Data represent means ± SEM of LC3 area per cell from 5 different photographic fields per mouse. **(C)** LC3 and LAMP2A *en face* staining of the aorta of *HDAC6*^*-/-*^*/ApoE*^*-/-*^ mice and littermate controls *HDAC6*^*+/+*^*/ApoE*^*-/-*^; n=6 per group; (red, LC3; green, LAMP2A; blue, DAPI). **(D)** Data represent means ± SEM of the percentage of LC3 and LAMP2A colocalization from 6 mice per group; \**P* ≤ 0.05 (Mann-Whitney test).

### Deletion of *HDAC6* reduces atherosclerotic lesions in hypercholesterolemic mice

To evaluate the effect of HDAC6 deletion on atherosclerotic plaque development, we assessed the plaque size by Oil Red O staining. Before initiating the high fat diet, both groups of mice were transplanted with *HDAC6*^*+/+*^*/ ApoE*^*-/-*^ bone marrow in order to rule out the effects of HDAC6 deletion in immune cells and mainly focus on its role on the vessel wall [35]–[38].

The high fat regimen resulted in a doubling of plasma cholesterol levels (Supplementary Figure 8A). *HDAC6* knockout, in ApoE^-/-^ mice transplanted with *HDAC6*^*+/+*^*/ ApoE*^*-/-*^ bone marrow, had no effect on cholesterol levels (Supplementary Figure 8A), body and organ weight, nor did it alter cell count for different populations in the blood (Supplementary Figure 8B) or the immune cell profile in the spleen (Supplementary Figure 8C) and bone marrow (Supplementary Figure 8D). As previously, described, atherosclerotic plaques preferentially formed low SS areas such as the inner curvature of the aortic arch and around carotids, whereas high SS areas, such as the descending aorta, remained relatively protected (Figure 8A). We found that plaque size was reduced by approximately 30% in the aortic arch of *HDAC6*-knockout animals when compared to littermate controls, whereas no significant differences were observed in the descending aortas (Figure 8B).

**Figure 8.**
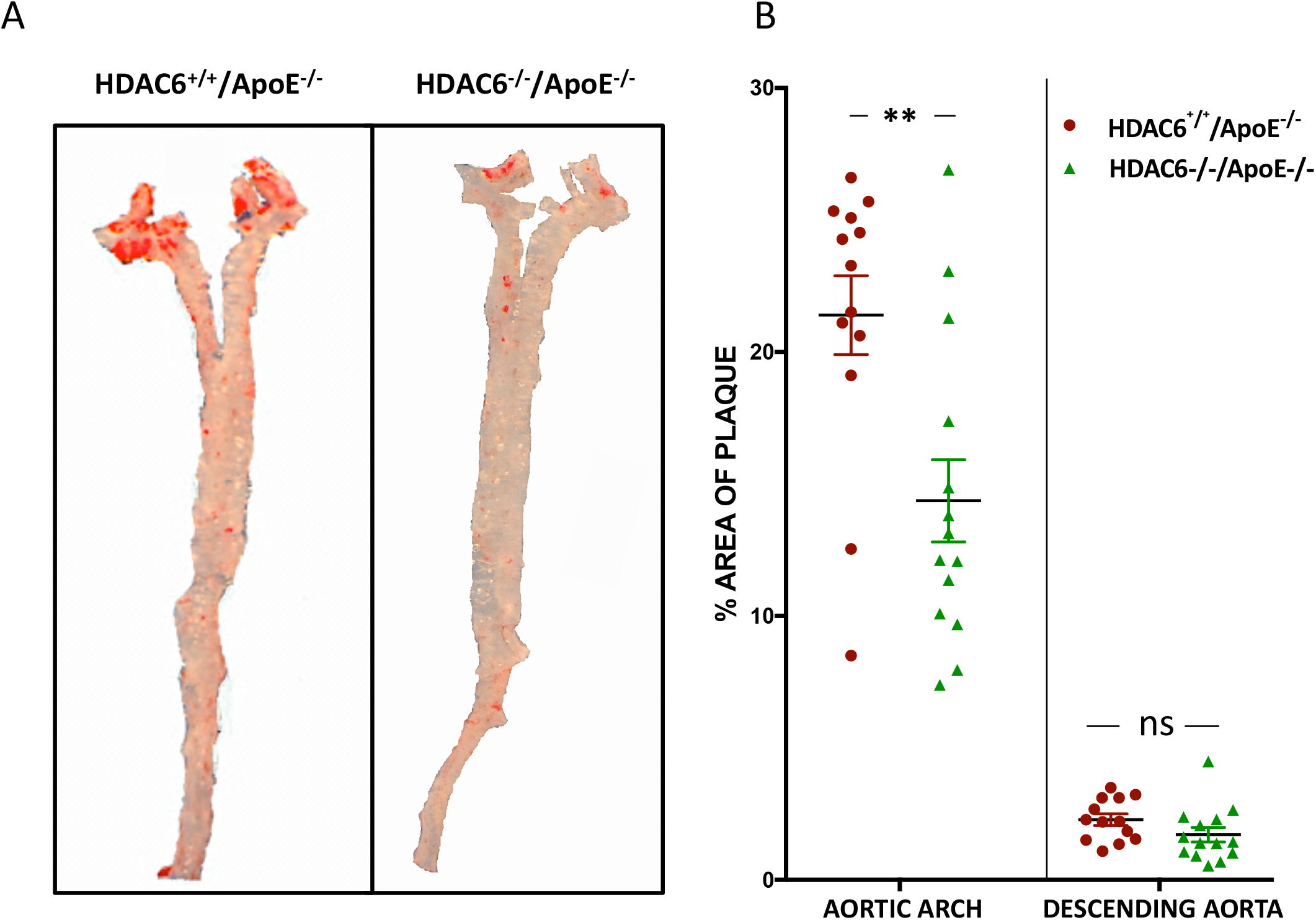

## DISCUSSION

Defects in endothelial autophagy occur in low shear stress areas of the vasculature, thereby favoring inflammation and the development of atherosclerotic lesions [6]. The present study demonstrates that restoration of adequate levels of endothelial autophagic flux, via HDAC6-inhibition, reduces endothelial inflammation and atherosclerotic plaque development, despite unfavorable shear stress conditions.

Atheroprone shear stress increases cytosolic HDAC6 activity, as demonstrated by the greater Ser22 HDAC6 phosphorylation in the endothelial cytosol of cells exposed to low SS. Interestingly other pro-atherogenic stimuli, such as cigarette smoke, also increase HDAC6 phosphorylation [39]. Upstream kinases of HDAC6 phosphorylation may include glycogen synthase kinase 3β (GSK-3β), which is activated by fluid flow through a Gα_q/11_-Akt-1-dependent pathway [19]. Moreover, HDAC6 activation may also be relayed through the zeta isoform of Protein Kinase C, which is stimulated by atheroprone shear stress [40]. We also observed that increased HDAC6 activation under low shear stress associates with decreased endothelial α-tubulin acetylation on lysine 40, indicative of lower microtubule stability [19], [41]. Under these conditions, endothelial α-tubulin acetylation was augmented after silencing HDAC6 mRNA or using the HDAC6 inhibitor Tubastatin-A, which has a 1000-fold greater specificity for this isoform than for other members of the HDAC superfamily [28], [42].

Using the LC3-RGP-GFP tandem, we found that Tubastatin-A restored *in vitro* endothelial autophagic flux in cells exposed to low SS conditions, reaching levels that were not different from cells under high SS, despite unfavorable shear stress conditions. Similarly, we observed a greater colocalization of LC3 and the lysosomal maker LAMP-2, in the aortic arch of *HDAC6*^-/-^ mice when compared to littermate controls, indicating a more robust *in vivo* autophagic flux in endothelial cells lacking HDAC6 and exposed to low SS. Interestingly, silencing HDAC6 *in vivo* resulted in comparable levels of endothelial autophagic flux in atheroprone (aortic arch) and atheroprotected (descending aorta) areas of the vasculature. Furthermore, our *in vitro* experiments performed in the presence of chloroquine revealed that HDAC6 inhibition also increased the rate of autophagosome biogenesis, without modifying the expression levels of essential autophagy proteins such as ATG5.

Several mechanisms could contribute to the beneficial effect of HDAC6 inhibition on autophagic flux. One possible interpretation is that increased autophagosome formation following HDAC6 inhibition might result, at least in-part, from increased trafficking of autophagic structures along acetylated microtubules. Hyperacetylation of tubulin enhances the recruitment of motor proteins to microtubules, which subsequently contribute to the concentration of signaling pathways involved in autophagosome biogenesis and to the transport of mature autophagosomes towards lysosomes [43], [44]. Several studies have indeed described impaired autophagosome and autolysosome formation in conditions where microtubules were disassembled using taxol or nocadazol [18], [43], [45]. More recently, Majora et al. demonstrated that Tubastatin-A and the pan-HDAC inhibitor, suberoylanilide hydroxamic acid increased tubulin acetylation and consequently improved autophagic function in human fibroblasts, in the context of Cockayne syndrome [46]. Another possible interpretation is that the augmented autophagy flux caused by endothelial HDAC6 inhibition results from increased expression and nuclear translocation of endothelial TFEB we report in low SS conditions, and which corroborates previous findings in experimental kidney disease [27]. Exposure of endothelial cells to high SS conditions activates TFEB, and TFEB endothelial overexpression increases lysosomal biogenesis and autophagic flux [24], [25]. However, the present study contrasts with these findings. Indeed, increased endothelial TFEB activity caused by HDAC6 inhibition or silencing is not associated with significant increased expression of two major lysosomal components LAMP1 and LAMP2A, despite increased LAMP2A-LC3 colocalization and augmented autophagolysosome formation as shown with the RFP-GFP-LC3 tandem experiments. The discrepancy between the different studies may results from differences in experimental settings compared with previous studies.

We then examined the functional consequences of endothelial autophagic flux improvement in atheroprone conditions, in particular whether or not HDAC6 inhibition or deficiency would affect endothelial inflammation and atherosclerotic plaque formation [6]. *In vitro* inhibition or silencing of HDAC6 reduced levels of reactive oxygen species and active NF-κB as well as the expression of inflammatory markers (ICAM-1, VCAM-1 and MCP-1) in endothelial cells exposed to low SS. These findings are in accordance with previous work showing an anti-inflammatory effect of Tubastatin-A in epithelial cells, *via* the regulation of the NF-κB pathway [47]. Other studies have shown that HDAC6 inhibition downregulates the production of inflammatory cytokines, such as IL-6, IL-1β, and TNF-α in murine models of arthritis and synovial inflammation, and Freund’s complete adjuvant-induced mouse model of inflammation [48]. HDAC6 inhibition has also been reported to reduce TNF-α-induced endothelial dysfunction and prolong survival in murine models of systemic inflammation and injury [49]. The molecular mechanisms underlying the anti-inflammatory effect of endothelial HDAC6 inhibition might be numerous. First, the autophagic machinery likely plays a central role in endothelial cells. Indeed, reducing levels of the essential autophagy protein ATG5 significantly reduced LC3-lipidation and suppressed the anti-inflammatory effects of HDAC6-inhibition by approximately 70%. This indicates that a functional autophagic process is key to relaying these beneficial effects; this interpretation is in line with our previous data showing that autophagy is required for atheroprotection under physiological blood flow [6]. Efficient autophagy may favor an anti-inflammatory endothelial phenotype by limiting the accumulation of damaged organelles, protein aggregates or intracellular pathogens [50], [51] and by limiting NF-κB signaling [52]. Second, the increased TFEB expression and nuclear translocation observed in the present study could also contribute to the anti-inflammatory effects of HDAC6 inhibition in low SS conditions, since TFEB overexpression decreases reactive oxygen species formation, NF-κB activation and impairs endothelial inflammation [25]. However, the increase in TFEB activity does not fully explain the anti-inflammatory effects of endothelial HDAC6 inhibition or silencing we observed in vitro. Indeed, endothelial TFEB overexpression has potent anti-inflammatory effects that are insensitive to autophagy inhibition (Lu et al, 2017), whereas the anti-inflammatory effects of HDAC6 inhibition or silencing observed in the present study are strongly impaired when ATG5 expression is inhibited. Therefore, it is plausible that TFEB contributes to a small part, if any, of anti-inflammatory effects resulting from HDAC6 inhibition or silencing.

To further assess the role of HDAC6 in atherosclerotic plaque formation, we used a double knockout chimeric murine model, *HDAC6*^*-/-*^*/ApoE*^*-/-*^. As previously described *HDAC6*^*-/-*^ animals were viable and showed no signs of major defects [53]. These mice, and their control littermates, underwent bone marrow transplantation, which as mentioned above, allowed us to focus on the role played by resident cells in the vasculature in the development of atherosclerotic plaques, whilst eliminating the effect of HDAC6 deletion in immune cells. We found that knocking out *HDAC6* reduced atherosclerotic plaque development in the low SS areas of hypercholesterolemic mice. These effects were not caused by other systemic metabolic parameters influencing cardiovascular risk, as plasma cholesterol levels, body weight and blood cell count were not affected by *HDAC6* knockout and *HDAC6*^*+/+*^ */ApoE*^*-/-*^ bone marrow transplantation. In addition to clearing damaged intracellular material, restoring autophagic flux in endothelial cells exposed to low shear stress, may also aid in their ability to limit lipid accumulation and the subsequent inflammatory response [54], [55]. Alternatively, data by Manea et al. indicated that administering suberoylanilide hydroxamic acid to hypercholesterolemic mice reduced the progression of atherosclerotic lesions through the reduction of oxidative stress in the aorta [56]. Furthermore, Chen et al. (2019) found that another HDAC6 inhibitor (Tubacin) mitigates endothelial dysfunction by regulating expression of endothelial nitric oxide synthetase [57]. Whether or not these alternate pathways are linked to the autophagic machinery would requires further investigation. While our focus in this study has been on endothelial cells, we cannot exclude that autophagy stimulation may also occur in smooth muscle cells via *HDAC6*-knockout. This very well might contribute to the anti-atherogenic effects we observed in our *in vivo* experimental model, since defective autophagy in vascular smooth muscle cells has been linked to accelerated atherogenesis [58].

In conclusion, the main findings of this study show that targeting HDAC6 stimulates endothelial autophagic flux, decreases endothelial inflammation and impairs the formation of atherosclerotic lesions in mice. This bolsters the idea that autophagy-stimulating strategies might be beneficial in treating atherosclerosis, and present HDAC6-oriented interventions as an interesting therapeutic route.

## MATERIAL AND METHODS

### Human Umbilical Vein Endothelial cells (HUVECs)

HUVECs were obtained from 20 different donors (11 males, 9 females) (PromoCell; Heidelberg, Germany). Cells were plated at a density of 10,000 cells/cm^2^ and cultured in Endothelial Cell Basal Medium (ECBM, PromoCell; Heidelberg, Germany), supplemented with 2% Fetal Calf Serum (PromoCell; Heidelberg, Germany), growth factors (0.4% ECGS, 0.1 ng/mL EGF, 1 ng/mL ß-FGF), 90 µg/mL heparin, 1 µg/mL hydrocortisone, 10 µg/L Amphotericin B (Gibco™ -15290026), 100 IU/mL Streptomycin (Gibco™-15140148) and 100 IU/mL Penicillin (Gibco™-15140148). Confluent cells were detached with 0.025% Trypsin-EDTA (Gibco™ - 25300-054) in PBS (Gibco™ - 10010023) for 5 min at 37°C, followed by washing with complete medium and centrifuging at 600g for 10min.

### Lentiviral transduction

Lentiviruses expressing inducible shRNA (Sigma-Aldrich, MISSION™ shRNA inducible vectors) were used to silence HDAC and ATG5. Cells were infected with lentivirus carrying shHDAC6/shATG5 or shControl at MOI 5 for 24 hours in the presence of Hexamethidine Bromide (8 µg/mL; Sigma-Aldrich - H9268). Transduced cells were selected using puromycin (Sigma Aldrich #P9620) at 1 µg/mL. HUVECs were treated for about 10 days with Isopropyl β-D-1-thiogalactopyranoside (IPTG; Sigma-Aldrich - I6758) at 1mmol/mL.

### *In Vitro* shear stress system

Confluent HUVECs from passage 2-4 were cultured on glass slides coated with 0.2% gelatin, and placed in a parallel plate chamber perfused with circulating medium for 24 hours, under laminar flow as described earlier [6]. When specifically stated, cells were treated with Tubastatin-A (3 µM, Sigma-Aldrich), Rapamycin (1 µM; Sigma-Aldrich) or Chloroquine (Sigma-Aldrich C6628) and exposed for 24 hours under shear stress conditions. For control conditions, HUVECs were exposed to vehicle (Dimethyl Sulphoxide; 0.1 µL/mL, Sigma-Aldrich). For inflammation studies, the cells were treated with TNF-α (1 ng/mL, Peprotech) for the last 12 hours of the experiment.

### Western blot analysis

HUVECs were immediately washed twice with PBS and lysed in Radioimmunoprecipitation assay (RIPA) buffer consisting of 150 mmol/L NaCl, 50 mmol/L Tris HCl (pH=7.4), 2 mmol/L EDTA, 2 mmol/L activated orthovanadate, 0.5% deoxycholate, 0.2% Sodium Dodecyl Sulphate and supplemented with Complete^™^ Mini protease and phosphatase inhibitors cocktails (Roche, France). Then, the lysates were centrifuged for 15 minutes at 12500g, 4°C to remove cell debris. The supernatant was collected and stored at -80°C until further analysis. Protein concentration was estimated by Lowry’s protein assay method (DC™ Protein Assay, BioRad).

Protein samples were denatured using buffer containing Tris-carboxyethyl phosphine hydrochloride (20x XT-Reducing Agent, BioRad-1610792) and separated based on molecular weight by electrophoresis in 4-12% gradient gels (Criterion™, BioRad-3450123). Proteins were then transferred on either 0.45 µm nitrocellulose (BioRad) or Polyvinylidene Fluoride (PVDF) membranes; following which the membranes were incubated with Ponceau Red to verify the efficiency of the transfer process. Membranes were blocked with 5% (w/v) milk or Bovine Serum Albumin (BSA) in TBS supplemented with 0.1% Tween-20. To detect the protein of interest, membranes were then incubated with primary antibodies overnight at 4°C, with constant agitation (Supplementary Table 1). After three 10-minute washes, the membranes were incubated with secondary antibody coupled with Horseradish Peroxidase (anti-rabbit, anti-rat or anti-mouse, Amersham, GE Healthcare, 1/3000) for 1 hour at room temperature. Immunodetection was performed using Clarity™ Western ECL Substrate, and the chemiluminescent signal was revealed using the Las-4000 imaging system and quantified with MultiGauge software (Fujifilm, Japan)

### Immunofluorescence

Cells were fixed with either 100% ice-cold methanol for 20 minutes at -20°C or 4% (v/v) paraformaldehyde (PFA) in PBS, for 5 minutes at room temperature. Cells fixed with PFA were permeabilized with 0.1% Triton X-100 for 10 minutes at room temperature. Permeabilized cells were blocked with 5% (w/v) BSA in PBS for 30 minutes and then incubated with primary antibody overnight at 4°C (Supplementary Table 2). The cells were washed with PBS and incubated with secondary antibody conjugated with fluorescent dye, Alexa Fluor^®^. Lastly, the nuclei were labeled with DAPI. The labeled cells were mounted on coverslips using Fluoromount-G^®^.

To perform *en face* immunostaining, aortas were harvested from mice and fixed with ice-cold 4% (v/v) PFA for 20 minutes. Residual fat attached to the aorta was removed and dissected along the entire inner side and outer curvature of the arch. Aortas were then blocked with 5% (w/v) BSA in PBS, for 60 minutes at room temperature, and incubated with primary antibody for overnight, 4°C. Following incubation with primary antibody, the aortas were washed multiple times and incubated with secondary antibody conjugated with Alexa Fluor^®^ for 2 hours at room temperature. Subsequently, the aortas were washed with PBS and the nuclei were labeled with DAPI. Labeled aortas were transferred onto glass slides and mounted “*en face*”. The slides prepared were examined using Leica SP8 confocal microscope and analyzed with software ImageJ. For colocalization analysis, ImageJ PlugIn *‘Colocalization Finder’* was used and Manders’ coefficients were calculated.

### Deacetylase activity assay

To determine the deacetylase enzymatic activity of HDAC6, a fluorescence-based assay was performed [59]. Deacetylation of substrate by HDAC6 sensitizes the substrate to the developer solution, which generates a fluorophore. The fluorescence signal is then detected by a fluorimetric plate reader at 460 nm. HUVECs were lysed in PBS buffer supplemented with - 0.5% Triton X-100, 300 mM NaCl and protease/phosphatase inhibitor cocktail (Thermo Fisher). Lysed cells were sonicated and centrifuged at 12500g for 15 minutes, 4°C. The pellet was discarded, and the supernatant was transferred to fresh tubes. Protein concentration was estimated by Micro BCA assay (Thermo Scientific™ - 23235). For the assay, 15 µg of lysates were diluted in 100 µL of PBS and added to a 96-well plate. Where required, 1 µL of the 100x HDAC6 inhibitor of the stock solution were added. Vehicle (DMSO) was added to the controls at the same concentration. The plates were incubated at 37°C for 60 minutes. 5 µL of substrate, specific to HDAC Class II, were added to the wells at a concentration of 1mM and incubated for 2 hours at 37°C. Substrate was bought from Bachem (I-1875; Boc-Lys (Ac)-AMC). 50 µL per well, developer solution was added and incubated at 37°C for 20 minutes. The developer solution consisted of 1.5% Triton X-100, 3 μM TSA (Sigma-Aldrich, T8552), and 0.75 mg/mL trypsin (Gibco™ 15400054), diluted in 1x PBS. AMC fluorescence was measured using a plate reader, with excitation and emission filters of 360 nm and 460 nm, respectively. Background signals corresponding to the blank solutions was subtracted.

### Monitoring autophagic flux using mRFP-GFP-LC3

In order to observe the effect of shear stress on endothelial autophagic flux, we used a LC3 protein, fluorescently tagged with a RFP-GFP tandem, (generous gift from Dr F. Oury, Inserm Paris), according to Vion, Kheloufi et al, [6]. The tandem allows us to observe different stages autophagy. At the autophagosomes stage, the LC3 protein is visualized as yellow signal, due to the overlapping of green and red fluorescence signal from GFP and RFP. However, when autolysosomes are formed, the acidic pH from lysosomes causes quenching of GFP. Therefore, the red fluorescent signal corresponds to LC3 present only in the autolysosomes stage. The cells were infected with lentiviruses carrying the plasmid (MOI = 6) in the presence of Hexamethidine Bromide (8 µg/mL; Sigma-H9268) for 24 hours. HUVECs were then exposed to shear stress for 24 hours with or without Tubastatin-A (3 µM; Sigma-SML0044). For experiments where autophagic flux was blocked, HUVECs were treated with Chloroquine (300µM) for the last 6 hours of the shear stress. After exposure to shear stress, the cells were washed with PBS and fixed with 4% PFA for 5 minutes at room temperature, followed by labeling the nuclei with DAPI. The slides were mounted using Fluoromount-G^®^ (SouthernBiotech). Images of the prepared slides were taken with the Leica SP8 confocal microscope and analyzed for LC3 positive signal using ImageJ software.

### ELISA

Levels of Monocyte Chemoattractant Protein-1 (MCP-1), released by HUVECs, in the media, were assessed by ELISA (R&D, Human MCP-1 Duo-Set DY279). The assay was performed according to manufacturer’s protocol.

### ROS Measurement

To assess ROS levels following exposure to shear stress, we used the CellROX Deep Red Reagent (Thermo Fisher Scientific, C10422); a fluorogenic probe designed to reliably measure ROS inside living cells. This cell-permeable dye is in a non-fluorescent reduced state while outside the cell and exhibits excitation/emission maxima at 640/665 nm upon oxidation. After 24h of shear stress exposure, HUVECs were incubated with 5 μmol/L CellROX for 30 minutes at 37°C. Cells were the then detached and analyzed using the LSRFortessa™ flow cytometer.

### Animal model

All the mice used in the study were of C57BL/6 genetic background. The double-knockout model was generated by crossing *HDAC6*^*-/-*^ mice, provided by T. McKinsey, University Colorado Denver, USA, and *ApoE*^*-/-*^ (Charles River Laboratories). All experiments were performed in accordance with the European Community guidelines for the care and use of laboratory animals (no. 07430) and were approved by the institutional ethical committee (no. 02526.02). All experiments were performed on male mice. Since the *HDAC6* gene is found on the X-chromosome, it was possible to obtain only male littermate control mice to be used for the experiments.

### Bone Marrow Transplant

8 weeks old *HDAC6*^*-/-*^*/ApoE*^*-/-*^ and *HDAC6*^*+/+*^*/ApoE*^*-/-*^ male mice underwent Bone Marrow Transplant (BMT) procedure, in order to replenish the bone marrow in the mutated models with *HDAC6*^*+/+*^*/ApoE*^*-/-*^ donor bone marrow. This allowed us to focus on the role of endothelial cells and rule out the potential function of hematopoietic cells in progression of atherosclerosis. 4-6 weeks old *HDAC6*^*+/+*^*/ApoE*^*-/-*^ donor mice were euthanized by cervical dislocation and the bone marrow was extracted from tibia and femur. The bone marrow was flushed and filtered. The recipient mice received radiation, 9.5 Gray, and were transplanted with donor bone marrow by retro-orbital injections the day after. The mice were allowed to recover for 4 weeks, following which they were put on high fat diet (D12079B, Research Diets; 20% proteins, 50% glucose, 21% lipids). After 10 weeks of high fat diet, the mice were euthanized.

### Evaluation of Blood Cholesterol Levels

Mice were anesthetized with 2% isoflurane and blood (75 µL) was collected by making a sub-mandibular puncture. To prepare platelet-free plasma, the blood collected was centrifuged twice at 2500g for 15 minutes at room temperature. The cholesterol content was determined using Cholesterol FS 10’ kit (Diasys).

### Isolation of Aortas

Mice were anesthetized using isoflurane (flow was set at 2.5% for induction and 2% for maintenance) and 2L/min of O_2_. The abdominal organs were moved aside to expose the inferior *vena cava*. 500-µL blood was drawn from the inferior *vena cava* using a 1 mL syringe and 26G needle primed with sodium citrate. Following blood collection, the diaphragm was cut, and the thoracic cavity exposed. The right atrium was nicked to release the blood. The heart was flushed with PBS via the left ventricle, using a 10 mL syringe and 25G needle. Followed by PBS, the heart was flushed with 1-2 mL ice-cold 4% PFA and PBS to wash out the excess PFA. To allow for better access to the aorta, the liver, lungs, trachea, oesophagus, thymus gland and pulmonary artery, were removed. Using micro-dissection scissors and forceps the excess adipose tissue and adventitia, surrounding the thoracic aorta and the carotids, were removed. The clean aorta was excised from the heart and placed in 4% PFA for 20 minutes on ice, followed by storage in ice-cold PBS.

### Determination of Atherosclerotic Plaque Size On *En Face* Aortas

The aortas were then stained with freshly prepared Oil Red O dye (40% distilled water, 60% Oil Red O solution) for 20 minutes with constant agitation. The solution was prepared by dissolving the dye in isopropanol at a concentration of 5g/L. The aortas were washed with 75% ethanol for 5 minutes to remove the excess stain and mounted “e*n face*” on glass slides and observed under microscope. Surface area occupied by the plaque was assessed using software, ImageJ.

### Preparation of Spleen

Spleens were then weighed and rinsed twice with ice-cold 0.1 μm filtered PBS containing 3% FBS. Placing them on 3% PBS-FBS pre-wetted 40 μm nylon cell strainers (Cat#352340, Thermo Fisher Scientific, USA) suspended over a 50 mL polypropylene tube, the spleens were then mashed using the plunger end of a 1 mL syringe. The cell strainers were then rinsed four times with 1 mL of 3% PBS-FBS. Cell suspension was then centrifuged at 500xg for 10 minutes at 4°C. The supernatant was then carefully aspirated and discarded, and the pellet was then resuspended in 1 mL of 0.1 μm filtered PBS. 1 mL of red blood cell lysis buffer (Cat# R7757, Merck, USA) was added to the resuspension and incubated at 5 minutes at room temperature. After the incubation, 18 mL of 3% PBS-FBS was added and suspension was centrifuged at 500g for 10 minutes at 4°C. Supernatant was then removed carefully, and pellet was resuspended in 3 mL of complete RPMI media that consisted of RPMI 1640 Medium with GlutaMAX and HEPES (Cat# 72400021, Thermo Fisher Scientific, USA) supplemented with 10% FBS, 1% P/S and 0.1% β-mercaptoethanol (Cat# 1610710, Bio-Rad, USA). Cells were then counted, and a final concentration of 10 × 10 ^6^ cells/mL was obtained.

### Preparation of Bone Marrow

The femur and tibia were isolated from the hind legs of the mouse. Sterilized micronic tubes, with a pore at the bottom, were placed in 1.5 mL Eppendorf tubes. One bone was placed per tube and centrifuged at 10000g for 15 seconds at room temperature. Collected cells were resuspended in 800 µL PBS, passed through 40micron syringe filter. The samples were then centrifuged at 400g for 10 minutes at 4°C. The pellet was resuspended in PBS to have a concentration of 10*10^6^ cells/mL. Bone marrow cells were incubated with the antibody mix for 30 minutes at 4°C. The plate was centrifuged at 500g for 5 minutes at 4°C. The pellet was resuspended in 100 µL of 1% PBS-FBS.

### Flow Cytometry

Circulating and splenic immune cell populations were defined as follows: NK cells as NK1.1^+^, neutrophils as CD11b^+^/Ly6C^+^/Ly6G^+^, monocytes as CD11b^+^/Ly6C^+^/Ly6G^-^, dendritic cells as CD11b^+^/Ly6C^+^/Ly6G^-^/CD11c^+^, macrophages as CD11b^lo/-^/Ly6c^+^/Ly6g^-^/F480^hi^, activated mature B lymphocytes as CD19^+^/IgM^+^/B220^+^, cytotoxic T lymphocytes as CD3^+^/CD8^+^ and helper T lymphocytes as CD3^+^/CD4^+^. Antibody incubation was performed on ice for 30 minutes (Supplemental Table 3) in 96-well plate and 100 µl of PBS-FBS 1% was added post-incubation. The plate was then centrifuged at 500g for 5 minutes at 4°C and supernatant was removed by inverting the plate. A final resuspension of splenocyte pellet with 100 µl of PBS-FBS 1% was performed before transfer to Micronic tubes (Cat# 2517810, Dutscher, France) for flow cytometry analysis (LSRFortessa™, BD Biosciences, USA). Data was analyzed using FlowJo software (BD Biosciences, USA).

### Statistical Analysis

Data are expressed as mean ± SEM for all experiments. Comparisons between different SS conditions or between control and treatment conditions were performed by using a Wilcoxon test for *in vitro experiments*. Comparisons between groups of mice were performed by using the Mann–Whitney *U* test or 2-way Anova (with Sidak post-hoc test). Statistical analyses were performed using the GraphPad Prism 7 statistical package. All tests were two-sided and used a significance level of 0.05.

## Abbreviations

CQ: Chloroquine
ICAM-1: Intercellular adhesion molecule 1
HDAC6: Histone Deacetylase 6
HUVEC: Human umbilical vein endothelial cell
LC3: microtubule-associated protein 1 light chain 3 β
MCP-1: Monocyte chemoattractant protein 1
SS: Shear stress
TFEB: transcription factor EB
TNF-α: Tumor necrosis factor alpha
TUBA: Tubastatin-A
VCAM-1: Vascular cell adhesion protein 1

## Acknowledgement

The authors acknowledge Cecile Devue for her helpful contributions, Flow Cytometry Facility manager, Dr Camille Knosp, as well as members of the INSERM U970 Histology, Microscopy and ERI facilities.

## REFERENCES

[1] “Lee and Chiu - 2019 - Atherosclerosis and flow roles of epigenetic modu.pdf.” Accessed: Jul. 28, 2020. [Online]. Available: https://jbiomedsci.biomedcentral.com/track/pdf/10.1186/s12929-019-0551-8

[2] “Baeyens et al. - 2016 - Endothelial fluid shear stress sensing in vascular.pdf.” Accessed: Jun. 14, 2020. [Online]. Available: https://dm5migu4zj3pb.cloudfront.net/manuscripts/83000/83083/cache/83083.1-20160218150308-covered-253bed37ca4c1ab43d105aefdf7b5536.pdf

[3] “Souilhol et al. - 2020 - Endothelial responses to shear stress in atheroscl.pdf.”

[4] C. Souilhol et al., “Endothelial responses to shear stress in atherosclerosis: a novel role for developmental genes,” Nat. Rev. Cardiol., vol. 17, no. 1, pp. 52–63, Jan. 2020, doi: 10.1038/s41569-019-0239-5.

[5] J.-J. Chiu and S. Chien, “Effects of Disturbed Flow on Vascular Endothelium: Pathophysiological Basis and Clinical Perspectives,” Physiol. Rev., vol. 91, no. 1, pp. 327–387, Jan. 2011, doi: 10.1152/physrev.00047.2009.

[6] A.-C. Vion et al., “Autophagy is required for endothelial cell alignment and atheroprotection under physiological blood flow,” Proc. Natl. Acad. Sci., vol. 114, no. 41, pp. E8675–E8684, Oct. 2017, doi: 10.1073/pnas.1702223114.

[7] Y. Feng, D. He, Z. Yao, and D. J. Klionsky, “The machinery of macroautophagy,” Cell Res., vol. 24, no. 1, pp. 24–41, Jan. 2014, doi: 10.1038/cr.2013.168.

[8] X. Wen and D. J. Klionsky, “An overview of macroautophagy in yeast,” J. Mol. Biol., vol. 428, no. 9, pp. 1681–1699, May 2016, doi: 10.1016/j.jmb.2016.02.021.

[9] N. Mizushima, B. Levine, A. M. Cuervo, and D. J. Klionsky, “Autophagy fights disease through cellular self-digestion,” Nature, vol. 451, no. 7182, pp. 1069–1075, Feb. 2008, doi: 10.1038/nature06639.

[10] N. Mizushima, “A brief history of autophagy from cell biology to physiology and disease,” Nat. Cell Biol., vol. 20, no. 5, pp. 521–527, May 2018, doi: 10.1038/s41556-018-0092-5.

[11] J. M. M. Levy, C. G. Towers, and A. Thorburn, “Targeting autophagy in cancer,” Nat. Rev. Cancer, vol. 17, no. 9, pp. 528–542, Sep. 2017, doi: 10.1038/nrc.2017.53.

[12] J. Mialet-Perez and C. Vindis, “Autophagy in health and disease: focus on the cardiovascular system,” Essays Biochem., vol. 61, no. 6, pp. 721–732, Dec. 2017, doi: 10.1042/EBC20170022.

[13] A. S. L. Wong, Z. H. Cheung, and N. Y. Ip, “Molecular machinery of macroautophagy and its deregulation in diseases,” Biochim. Biophys. Acta BBA - Mol. Basis Dis., vol. 1812, no. 11, pp. 1490–1497, Nov. 2011, doi: 10.1016/j.bbadis.2011.07.005.

[14] F.-X. Guo, Y.-W. Hu, L. Zheng, and Q. Wang, “Shear Stress in Autophagy and Its Possible Mechanisms in the Process of Atherosclerosis,” DNA Cell Biol., vol. 36, no. 5, pp. 335–346, May 2017, doi: 10.1089/dna.2017.3649.

[15] J.-J. Chiu et al., “Shear Stress Increases ICAM-1 and Decreases VCAM-1 and E-selectin Expressions Induced by Tumor Necrosis Factor-α in Endothelial Cells,” Arterioscler. Thromb. Vasc. Biol., vol. 24, no. 1, pp. 73–79, Jan. 2004, doi: 10.1161/01.ATV.0000106321.63667.24.

[16] R. Mackeh, D. Perdiz, S. Lorin, P. Codogno, and C. Pous, “Autophagy and microtubules - new story, old players,” J. Cell Sci., vol. 126, no. 5, pp. 1071–1080, Mar. 2013, doi: 10.1242/jcs.115626.

[17] B. Ravikumar et al., “Dynein mutations impair autophagic clearance of aggregate-prone proteins,” Nat. Genet., vol. 37, no. 7, pp. 771–776, Jul. 2005, doi: 10.1038/ng1591.

[18] R. Xie, S. Nguyen, W. L. McKeehan, and L. Liu, “Acetylated microtubules are required for fusion of autophagosomes with lysosomes,” BMC Cell Biol., vol. 11, no. 1, p. 89, 2010, doi: 10.1186/1471-2121-11-89.

[19] S. McCue, D. Dajnowiec, F. Xu, M. Zhang, M. R. Jackson, and B. L. Langille, “Shear Stress Regulates Forward and Reverse Planar Cell Polarity of Vascular Endothelium In Vivo and In Vitro,” Circ. Res., vol. 98, no. 7, pp. 939–946, Apr. 2006, doi: 10.1161/01.RES.0000216595.15868.55.

[20] G. M. Birdsey et al., “The transcription factor Erg regulates expression of histone deacetylase 6 and multiple pathways involved in endothelial cell migration and angiogenesis,” Blood, vol. 119, no. 3, pp. 894–903, Jan. 2012, doi: 10.1182/blood-2011-04-350025.

[21] L. Eshun-Wilson et al., “Effects of α-tubulin acetylation on microtubule structure and stability,” Proc. Natl. Acad. Sci., vol. 116, no. 21, pp. 10366–10371, May 2019, doi: 10.1073/pnas.1900441116.

[22] K. Ustinova et al., “The disordered N-terminus of HDAC6 is a microtubule-binding domain critical for efficient tubulin deacetylation,” J. Biol. Chem., vol. 295, no. 9, pp. 2614–2628, Feb. 2020, doi: 10.1074/jbc.RA119.011243.

[23] S. Chen, G. C. Owens, H. Makarenkova, and D. B. Edelman, “HDAC6 Regulates Mitochondrial Transport in Hippocampal Neurons,” PLoS ONE, vol. 5, no. 5, p. e10848, May 2010, doi: 10.1371/journal.pone.0010848.

[24] W. Song et al., “Endothelial TFEB (Transcription Factor EB) Restrains IKK (IκB Kinase)-p65 Pathway to Attenuate Vascular Inflammation in Diabetic db/db Mice,” Arterioscler. Thromb. Vasc. Biol., vol. 39, no. 4, pp. 719–730, Apr. 2019, doi: 10.1161/ATVBAHA.119.312316.

[25] H. Lu et al., “TFEB inhibits endothelial cell inflammation and reduces atherosclerosis,” Sci. Signal., vol. 10, no. 464, Jan. 2017, doi: 10.1126/scisignal.aah4214.

[26] J. Zhang et al., “Importance of TFEB acetylation in control of its transcriptional activity and lysosomal function in response to histone deacetylase inhibitors,” Autophagy, vol. 14, no. 6, pp. 1043–1059, 2018, doi: 10.1080/15548627.2018.1447290.

[27] A. S. Brijmohan et al., “HDAC6 Inhibition Promotes Transcription Factor EB Activation and Is Protective in Experimental Kidney Disease,” Front. Pharmacol., vol. 9, 2018, doi: 10.3389/fphar.2018.00034.

[28] K. V. Butler, J. Kalin, C. Brochier, G. Vistoli, B. Langley, and A. P. Kozikowski, “Rational Design and Simple Chemistry Yield a Superior, Neuroprotective HDAC6 Inhibitor, Tubastatin A,” J. Am. Chem. Soc., vol. 132, no. 31, pp. 10842–10846, Aug. 2010, doi: 10.1021/ja102758v.

[29] “McCue et al. - 2006 - Shear Stress Regulates Forward and Reverse Planar .pdf.” Accessed: Jul. 28, 2020. [Online]. Available: https://www.ahajournals.org/doi/pdf/10.1161/01.RES.0000216595.15868.55

[30] M. Mauthe et al., “Chloroquine inhibits autophagic flux by decreasing autophagosome-lysosome fusion,” Autophagy, vol. 14, no. 8, pp. 1435–1455, 2018, doi: 10.1080/15548627.2018.1474314.

[31] C. Settembre et al., “TFEB Links Autophagy to Lysosomal Biogenesis,” Science, vol. 332, no. 6036, pp. 1429–1433, Jun. 2011, doi: 10.1126/science.1204592.

[32] G. Napolitano and A. Ballabio, “TFEB at a glance,” J. Cell Sci., vol. 129, no. 13, pp. 2475–2481, Jul. 2016, doi: 10.1242/jcs.146365.

[33] C. Di Malta, L. Cinque, and C. Settembre, “Transcriptional Regulation of Autophagy: Mechanisms and Diseases,” Front. Cell Dev. Biol., vol. 0, 2019, doi: 10.3389/fcell.2019.00114.

[34] Y. Zhang et al., “Rapamycin Promotes the Autophagic Degradation of Oxidized Low-Density Lipoprotein in Human Umbilical Vein Endothelial Cells,” J. Vasc. Res., vol. 52, no. 3, pp. 210–219, Dec. 2015, doi: 10.1159/000441143.

[35] P. J. Thul et al., “A subcellular map of the human proteome,” Science, vol. 356, no. 6340, May 2017, doi: 10.1126/science.aal3321.

[36] M. Uhlen et al., “Tissue-based map of the human proteome,” Science, vol. 347, no. 6220, pp. 1260419–1260419, Jan. 2015, doi: 10.1126/science.1260419.

[37] “The Human Protein Atlas.” https://www.proteinatlas.org/ (accessed Jul. 28, 2021).

[38] “The Human Protein Atlas.” Accessed: Jul. 28, 2021. [Online]. Available: https://www.proteinatlas.org/

[39] D. Borgas et al., “Cigarette Smoke Disrupted Lung Endothelial Barrier Integrity and Increased Susceptibility to Acute Lung Injury via Histone Deacetylase 6,” Am. J. Respir. Cell Mol. Biol., vol. 54, no. 5, pp. 683–696, 2016, doi: 10.1165/rcmb.2015-0149OC.

[40] Y. Du, M. L. Seibenhener, J. Yan, J. Jiang, and M. C. Wooten, “aPKC Phosphorylation of HDAC6 Results in Increased Deacetylation Activity,” PLoS ONE, vol. 10, no. 4, Apr. 2015, doi: 10.1371/journal.pone.0123191.

[41] K. A. Bailey, F. G. Haj, S. I. Simon, and A. G. Passerini, “Atherosusceptible Shear Stress Activates Endoplasmic Reticulum Stress to Promote Endothelial Inflammation,” Sci. Rep., vol. 7, no. 1, p. 8196, Dec. 2017, doi: 10.1038/s41598-017-08417-9.

[42] Y.-J. Choi, M.-H. Kang, K. Hong, and J.-H. Kim, “Tubastatin A inhibits HDAC and Sirtuin activity rather than being a HDAC6-specific inhibitor in mouse oocytes,” Aging, vol. 11, no. 6, pp. 1759–1777, Mar. 2019, doi: 10.18632/aging.101867.

[43] C. Geeraert et al., “Starvation-induced Hyperacetylation of Tubulin Is Required for the Stimulation of Autophagy by Nutrient Deprivation,” J. Biol. Chem., vol. 285, no. 31, pp. 24184–24194, Jul. 2010, doi: 10.1074/jbc.M109.091553.

[44] Á. Bánréti, M. Sass, and Y. Graba, “The emerging role of acetylation in the regulation of autophagy,” Autophagy, vol. 9, no. 6, pp. 819–829, Jun. 2013, doi: 10.4161/auto.23908.

[45] R. Köchl, X. W. Hu, E. Y. W. Chan, and S. A. Tooze, “Microtubules Facilitate Autophagosome Formation and Fusion of Autophagosomes with Endosomes: Role of Microtubules in AV Formation,” Traffic, vol. 7, no. 2, pp. 129–145, Feb. 2006, doi: 10.1111/j.1600-0854.2005.00368.x.

[46] M. Majora et al., “HDAC inhibition improves autophagic and lysosomal function to prevent loss of subcutaneous fat in a mouse model of Cockayne syndrome,” Sci. Transl. Med., vol. 10, no. 456, p. eaam7510, Aug. 2018, doi: 10.1126/scitranslmed.aam7510.

[47] J. Wang et al., “Inhibition of histone deacetylase reduces lipopolysaccharide-induced-inflammation in primary mammary epithelial cells by regulating ROS-NF-кB signaling pathways,” Int. Immunopharmacol., vol. 56, pp. 230–234, Mar. 2018, doi: 10.1016/j.intimp.2018.01.039.

[48] J. Ran and J. Zhou, “Targeted inhibition of histone deacetylase 6 in inflammatory diseases,” Thorac. Cancer, vol. 10, no. 3, pp. 405–412, Mar. 2019, doi: 10.1111/1759-7714.12974.

[49] J. Yu, M. Ma, Z. Ma, and J. Fu, “HDAC6 inhibition prevents TNF-α-induced caspase 3 activation in lung endothelial cell and maintains cell-cell junctions,” Oncotarget, vol. 7, no. 34, pp. 54714–54722, Jul. 2016, doi: 10.18632/oncotarget.10591.

[50] S. Mai, B. Muster, J. Bereiter-Hahn, and M. Jendrach, “Autophagy proteins LC3B, ATG5 and ATG12 participate in quality control after mitochondrial damage and influence lifespan,” Autophagy, vol. 8, no. 1, pp. 47–62, Jan. 2012, doi: 10.4161/auto.8.1.18174.

[51] K. Cadwell, “Crosstalk between autophagy and inflammatory signalling pathways: balancing defence and homeostasis,” Nat. Rev. Immunol., vol. 16, no. 11, pp. 661–675, 2016, doi: 10.1038/nri.2016.100.

[52] X. Peng et al., “ATG5-mediated autophagy suppresses NF-κB signaling to limit epithelial inflammatory response to kidney injury,” Cell Death Dis., vol. 10, no. 4, p. 253, Apr. 2019, doi: 10.1038/s41419-019-1483-7.

[53] Y. Zhang et al., “Mice Lacking Histone Deacetylase 6 Have Hyperacetylated Tubulin but Are Viable and Develop Normally,” Mol. Cell. Biol., vol. 28, no. 5, pp. 1688–1701, Mar. 2008, doi: 10.1128/MCB.01154-06.

[54] K. Torisu et al., “Intact endothelial autophagy is required to maintain vascular lipid homeostasis,” Aging Cell, vol. 15, no. 1, pp. 187–191, Feb. 2016, doi: 10.1111/acel.12423.

[55] H.-S. Kim et al., “Metformin reduces saturated fatty acid-induced lipid accumulation and inflammatory response by restoration of autophagic flux in endothelial cells,” Sci. Rep., vol. 10, no. 1, Art. no. 1, Aug. 2020, doi: 10.1038/s41598-020-70347-w.

[56] S.-A. Manea et al., “Pharmacological inhibition of histone deacetylase reduces NADPH oxidase expression, oxidative stress and the progression of atherosclerotic lesions in hypercholesterolemic apolipoprotein E-deficient mice; potential implications for human atherosclerosis,” Redox Biol., vol. 28, p. 101338, Jan. 2020, doi: 10.1016/j.redox.2019.101338.

[57] J. Chen et al., “The histone deacetylase inhibitor tubacin mitigates endothelial dysfunction by up-regulating the expression of endothelial nitric oxide synthase,” J. Biol. Chem., vol. 294, no. 51, pp. 19565–19576, Dec. 2019, doi: 10.1074/jbc.RA119.011317.

[58] M. O. Grootaert et al., “Defective autophagy in vascular smooth muscle cells accelerates senescence and promotes neointima formation and atherogenesis,” Autophagy, vol. 11, no. 11, pp. 2014–2032, Nov. 2015, doi: 10.1080/15548627.2015.1096485.

[59] D. D. Lemon et al., “Cardiac HDAC6 catalytic activity is induced in response to chronic hypertension,” J. Mol. Cell. Cardiol., vol. 51, no. 1, pp. 41–50, Jul. 2011, doi: 10.1016/j.yjmcc.2011.04.005.

