## Supplemental Table 1 for "Restoring vascular endothelial autophagic flux reduces atherosclerotic lesions"

| **ANTIBODY** | **HOST SPECIES** | **REF** | **DILUTION** | **DILUTION**  **BUFFER** |
| --- | --- | --- | --- | --- |
| **LC3-B** | Rabbit | CST-2775S | 1/1000 | 5% milk +TBST |
| **GAPDH** | Mouse | mab-374 | 1/20000 | 5% milk +TBST |
| **ATG5** | Rabbit | CST-12994S | 1/1000 | 5% milk +TBST |
| **ATG7** | Rabbit | CST-2631S | 1/1000 | 5% milk +TBST |
| **p62** | Mouse | ab56162 | 1/3000 | 5% milk +TBST |
| **Beclin-1** | Rabbit | CST-3495S | 1/1000 | 5% milk +TBST |
| **HDAC6** | Rabbit | CST-7558S (D2E5) | 1/1000 | 5% milk +TBST |
| **phospho-HDAC6** | Mouse | ab61058 | 1/3000 | 5% BSA+TBST |
| **Acetylated α-Tubulin** | Mouse | Sigma T7451 | 1/3000 | 5% milk +TBST |
| **Total α-Tubulin** | Rat | ab6160 | 1/5000 | 5% milk +TBST |
| **ICAM-1** | Goat | R&D: AF796 | 1/1000 | 5% milk +TBST |
| **VCAM** | Rabbit | ab134047 | 1/3000 | 5% milk +TBST |
| **Phospho-p65** | Rabbit | CST-3033S | 1/1000 | 5% BSA+TBST |
| **TFEB** | Rabbit | **CST-4240S** | 1/1000 | 5% milk +TBST |
| **Phospho-TFEB** | Rabbit | **ABE1971-I** | 1/1000 | 5% BSA+TBST |

**Supplementary Table 1:** List of antibodies used for western blot analyses (TBST, Tris Buffer Saline 0.05%Tween)

| **ANTIBODY** | **HOST**  **SPECIES** | **REFERENCE** | **FIXATION** | **DILUTION** |
| --- | --- | --- | --- | --- |
| **LC3-B** | Rabbit | CST-2775S | 100% Methanol | 1/400 |
| **VE-Cadherin** | Goat | SCBT- (F-8) 9989 | -4% PFA | 1/400 |
| **HDAC6** | Rabbit | CST-7558S (D2E5) | 100% Methanol | 1/100 |
| **phospho-HDAC6 (Ser22)** | Mouse | ab61058 | 100% Methanol | 1/200 |
| **Acetylated α-Tubulin** | Mouse | Sigma T7451 | 4% PFA | 1/500 |
| **Total α-Tubulin** | Rat | ab6160 | 4% PFA | 1/1000 |

**Supplementary Table 2:** List of antibodies used immunofluorescence assays. Abbreviations: PBS, Phosphate Buffer Saline; PFA, Paraformaldehyde ; CST, Cell Signalling Technology, SCBT, Santa cruz Biotechnology ; ab,Abcam

| **ANTIBODY** | **FLUOROCHROME** | **REF** | **COMPANY** |
| --- | --- | --- | --- |
| Anti-NK1.1 | BV650 | 564143 | BD Biosciences |
| Anti-CD11b | FITC | 53-0112-82 | eBioscience |
| Anti-Ly6C | BV785 | 128041 | Biolegend |
| Anti-Ly6G | Pe-Dazzle594 | 127648 | Biolegend |
| Anti-CD11c | PerCP-Cy5.5 | 560584 | BD Biosciences |
| Anti-CD3e | BUV737 | 564618 | BD Biosciences |
| Anti-CD4 | BV510 | 563106 | BD Biosciences |
| Anti-CD8a | BV605 | 563152 | BD Biosciences |
| Anti-CD19 | Pe-Cy5 | 15-0193-83 | eBioscience |
| Anti-IgM | Pe-Cy7 | 406514 | Biolegend |
| Anti-CD45R/B220 | BV711 | 563892 | BD Biosciences |
| Anti-CD23 | PE | 12-0232-81 | eBioscience |
| Fixable Viability Dye | EF780 | 65-0865-14 | eBioscience |

**Supplementary Table 3:** List of antibodies used in flow cytometry experiments.

**Supplementary Figures**

Supplementary Figure 1

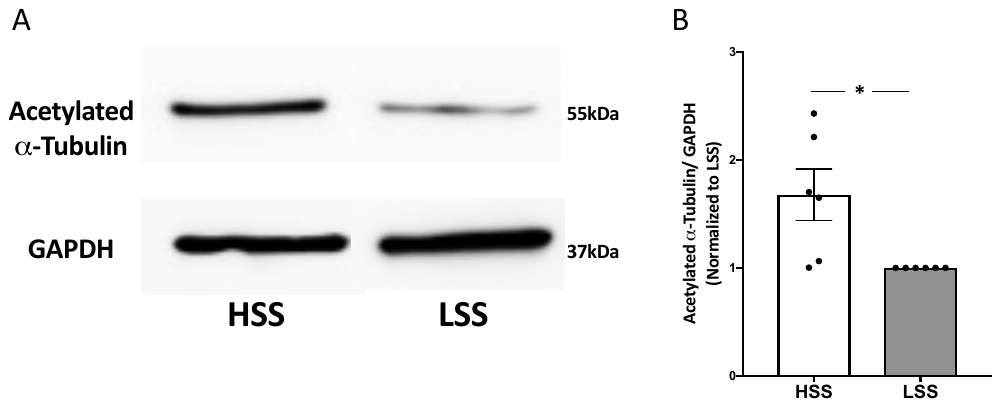

**Supplementary figure 1: Shear stress regulates acetylation of α-tubulin. (A)** Western blot analysis of acetylated α-tubulin expression in HUVECs exposed to either high SS (HSS; 20 dyn/cm^2^) or low SS (LSS; 2 dyn/cm^2^) for 24 hours. **(B)** Quantification of the ratio of acetylated α-tubulin to GAPDH normalized to low SS; data represent means ± SEM of 6 independent experiments. **P* ≤ 0.05 (Wilcoxon test).

Supplementary Figure 2

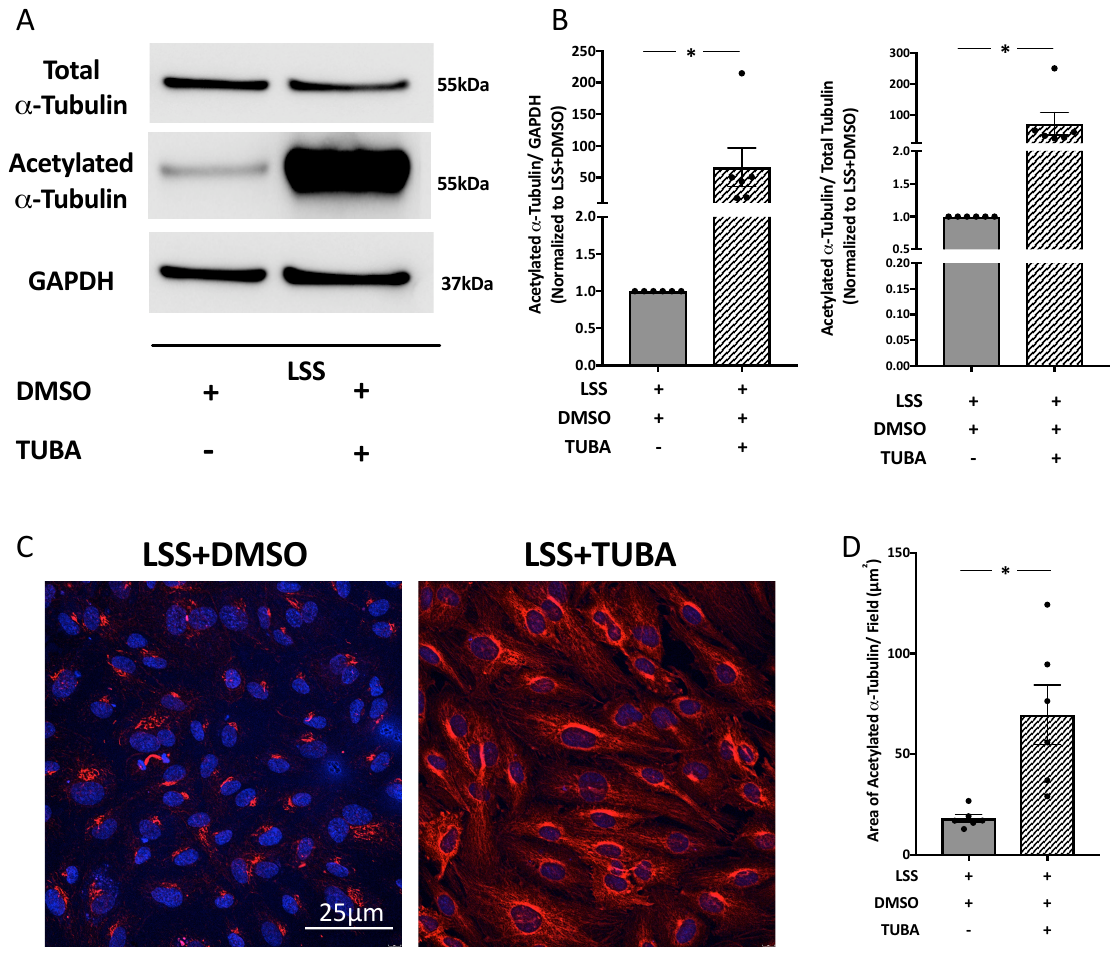

**Figure 2: Inhibition of HDAC6 activity by Tubastatin-A, increases levels of acetylated α-tubulin *in vitro*. (A)** Western blot analysis of acetylated α-tubulin in HUVECs exposed to low SS (LSS; 2 dyn/cm^2^) and treated with either vehicle (DMSO: 0.1 µL/mL) or Tubastatin-A (TUBA; 3 µM) for 24 hours. **(B)** Quantification of the ratio of acetylated α-tubulin to either GAPDH (left) or total tubulin (right); data represent means ± SEM of from 6 independent experiments, normalized to low SS conditions. **(C)** Representative confocal microscopy images of HUVECs exposed to low SS conditions for 24 hours and treated with or without Tubastatin-A (red: acetylated α-tubulin, blue: DAPI). **(D)** Data represent means ± SEM of 6 independent experiments. **P* ≤ 0.05 (Wilcoxon test).

Supplementary Figure 3

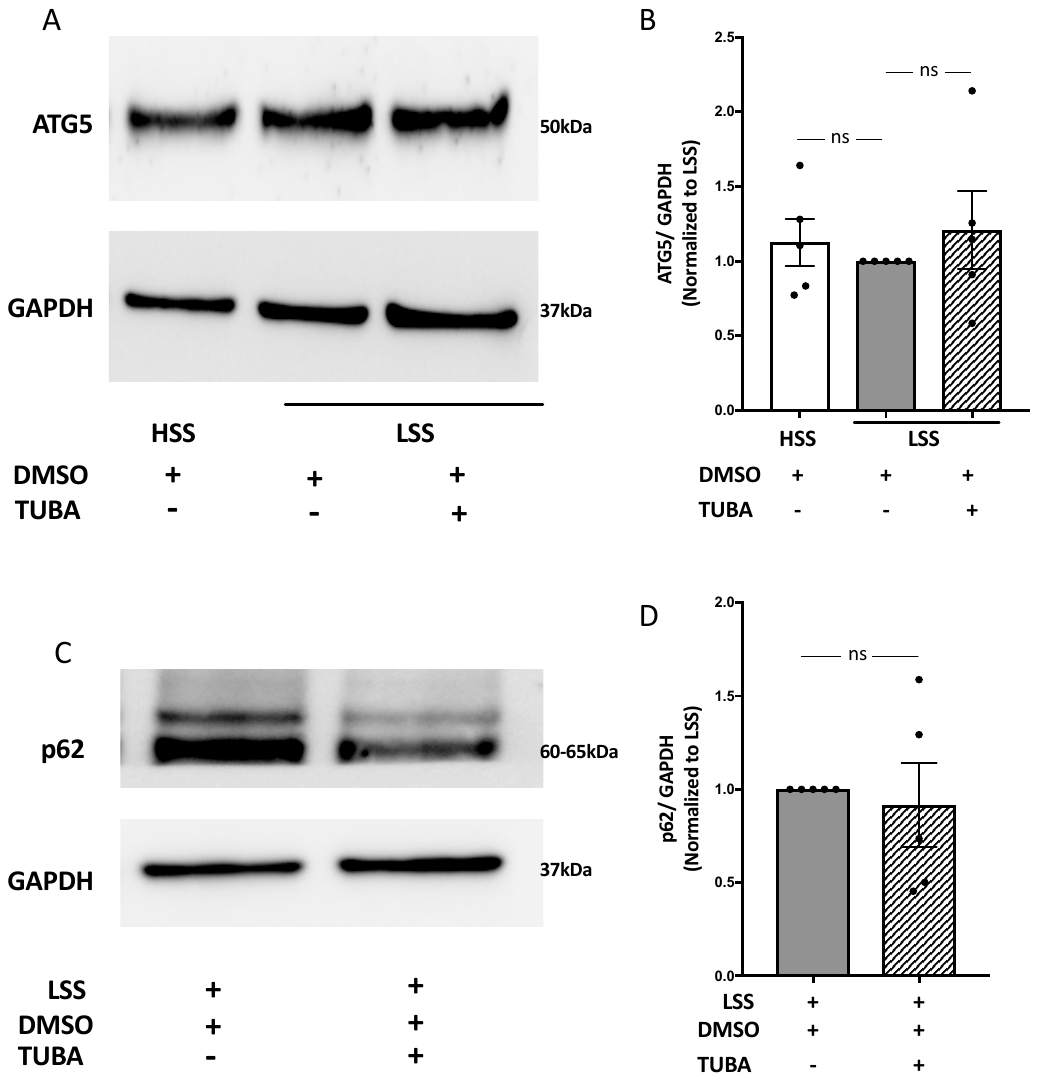

**Supplementary figure 3: Effect of Tubastatin-A on ATG5 and p62 expression. (A)** Western blot analysis of ATG5 expression in HUVECs exposed to high SS (HSS; 20 dyn/cm^2^) or low SS (LSS; 2 dyn/cm^2^) and treated with either vehicle (DMSO: 0.1 µL/mL) or Tubastatin-A (TUBA; 3 µM) for 24 hours. **(B)** Quantification of the ratio of ATG5 to GAPDH normalized to low SS + DMSO; data represent means ± SEM of 6 independent experiments. **(C)** Western blot analysis of p62 expression in HUVECs exposed to low SS (LSS; 2 dyn/cm^2^) and treated with either vehicle (DMSO: 0.1 µL/mL) or Tubastatin-A (TUBA; 3 µM) for 24 hours. **(D)** Quantification of the ratio of p62 to GAPDH normalized to low SS + DMSO; data represent means ± SEM of 6 independent experiments. ns, not statistically different (Wilcoxon test).

Supplementary Figure 4

**Supplementary figure 4: Effect of Tubastatin-A on TFEB expression.** Western blot analysis of **(A)** TFEB, LAMP1 AND **(C)** phosphorylated-TFEB expression in HUVECs exposed to high SS (HSS; 20 dyn/cm^2^) or low SS (LSS; 2 dyn/cm^2^) and treated with either vehicle (DMSO: 0.1 µL/mL) or Tubastatin-A (TUBA; 3 µM) for 24 hours. **(B)** Quantification of the ratio of phosphorylated-TFEB to total TFEB normalized to low SS + DMSO; data represent means ± SEM of 5 independent experiments. **(C)** Quantification of the ratio of LAMP1 to GAPDH and normalized to low SS + DMSO; data represent means ± SEM of 4 independent experiments. **(E)** Representative confocal microscopy images of HUVECs exposed to low SS conditions for 24 hours and treated with or without Tubastatin-A (red: TFEB, blue: DAPI). **(F)** Relative quantification of TFEB present in the **(G)** cytosol**/ (H)** nucleus. Data represent means ± SEM of 3 mice per group, with 5 different photographic fields per animal. **P* ≤ 0.05 (Two-way Anova, Sidak’s post-analysis test).

Supplementary Figure 5

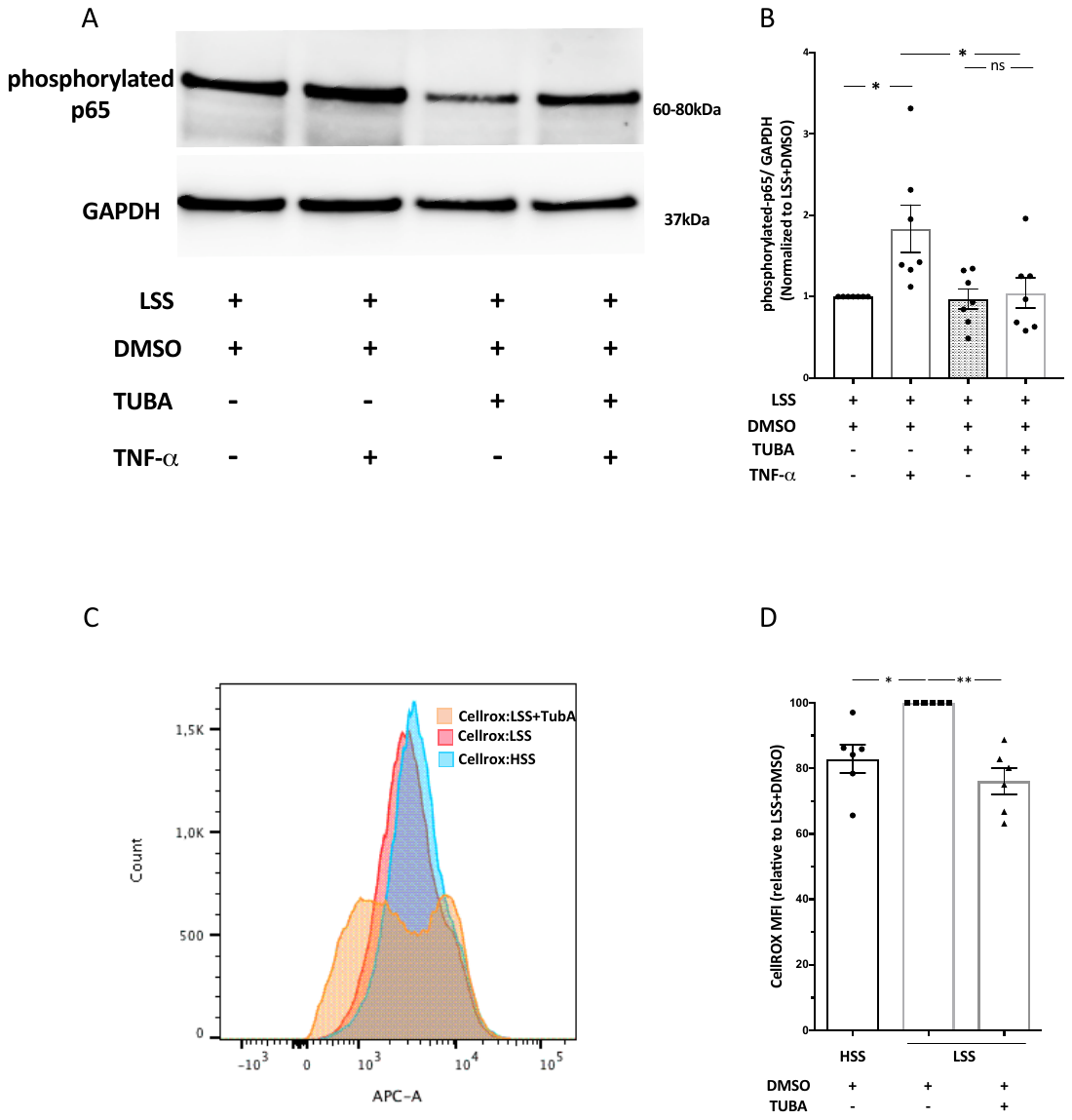

**Supplementary figure 5: Tubastatin-A blocks the increase of phospho-p65 in response to TNF-α and has anti-oxidative effect. (A)** Western blot analysis of p65 in HUVECs exposed to low SS (LSS; 2 dyn/cm^2^) and treated with either vehicle (DMSO at 0.1 µL/mL) or Tubastatin-A (TUBA; 3 µM) for 24 hours, in the presence or absence of TNF-α (1 ng/mL). **(B)** Quantification of the p65/GAPDH ratio; data represent means ± SEM of 4 independent experiments normalized to low SS + DMSO + TNF-α. **(C)** Representative plot and **(D)** Quantification of the ROS produced by HUVECs exposed to high SS treated with vehicle (HSS; 20 dyn/cm^2^); low SS (LSS; 2 dyn/cm^2^) and treated with either vehicle (DMSO at 0.1 µL/mL) or Tubastatin-A (TUBA; 3 µM) for 24 hours.

Supplementary Figure 6

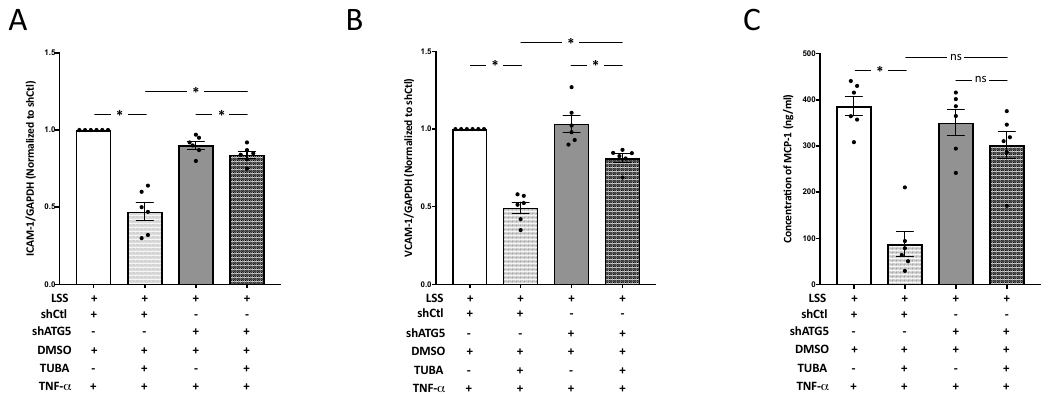

**Supplementary figure 6: Attenuation of Tubastatin-A’s anti-inflammatory effects in ATG5-deficient cells.** Analysis of ICAM-1 **(A)** and VCAM-1 **(B)** expression in endothelial cells transduced with either shControl (shCtl) or shATG5 lentiviruses, exposed to low SS (LSS; 2 dyn/cm^2^), and treated with vehicle (DMSO at 0.1 μL/mL) or Tubastatin-A (TUBA; 3 μM) in the presence of TNF-α (1 ng/mL). Data represent means ± SEM of 6 independent experiments normalized to shControl conditions treated with vehicle, DMSO. **(C)** ELISA analysis of MCP-1 levels released in the conditioned media of HUVECs. Data represent means ± SEM of 6 independent experiments. ns, not statistically different; **P* ≤ 0.05 (Wilcoxon test).

Supplementary Figure 7

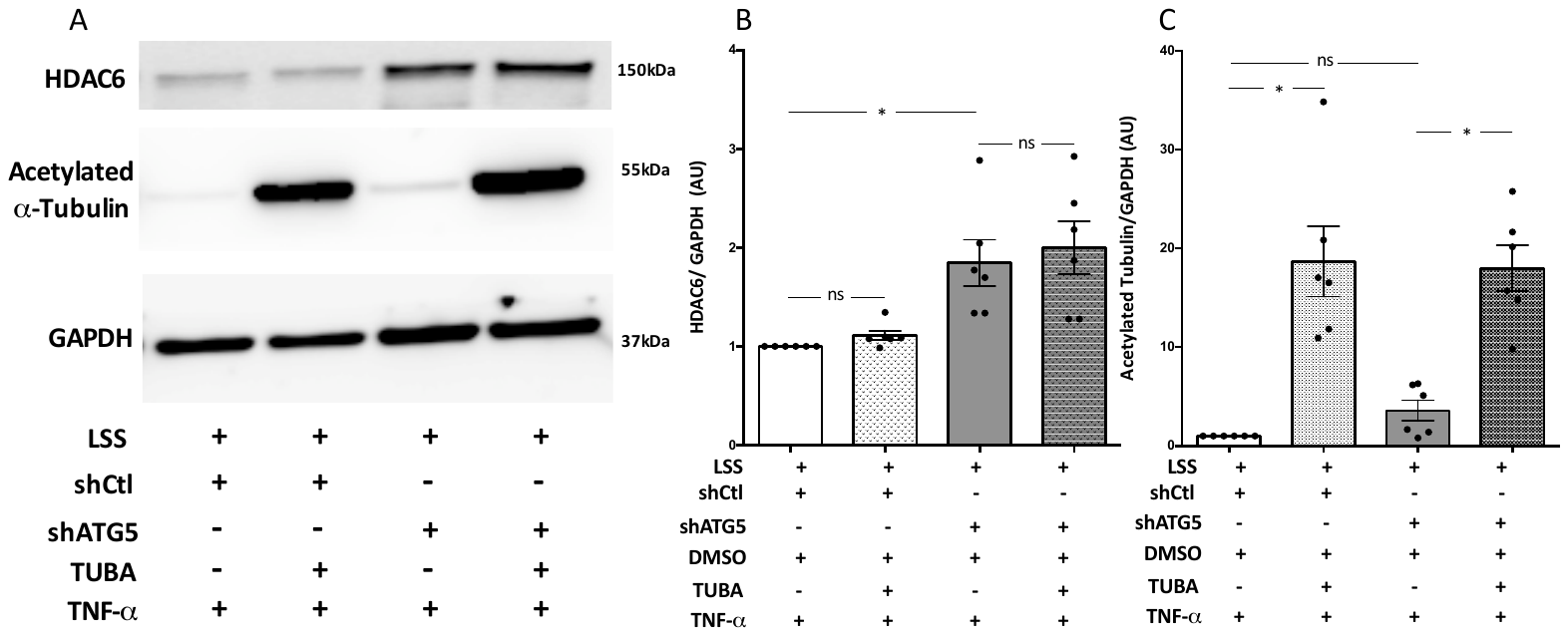

**Supplementary figure 7: Effect of ATG5 knockdown on expression of HDAC6 acetylated α-tubulin, under Tubastatin-A treated conditions: (A)** Western blots analysis of HDAC6 and acetylated α-tubulin in endothelial cells transduced with either shControl (shCtl) or shATG5 lentiviruses, exposed to low SS (LSS; 2 dyn/cm^2^), and treated with vehicle (DMSO at 0.1 μL/mL) or Tubastatin-A (TUBA; 3 μM) in the presence of TNF-α (1 ng/mL). Quantification of the HDAC6**/**GAPDH **(B)**, and Acetylated α-tubulin/GAPDH **(C)** ratios; data represent means ± SEM of 6 independent experiments.

Supplementary Figure 8

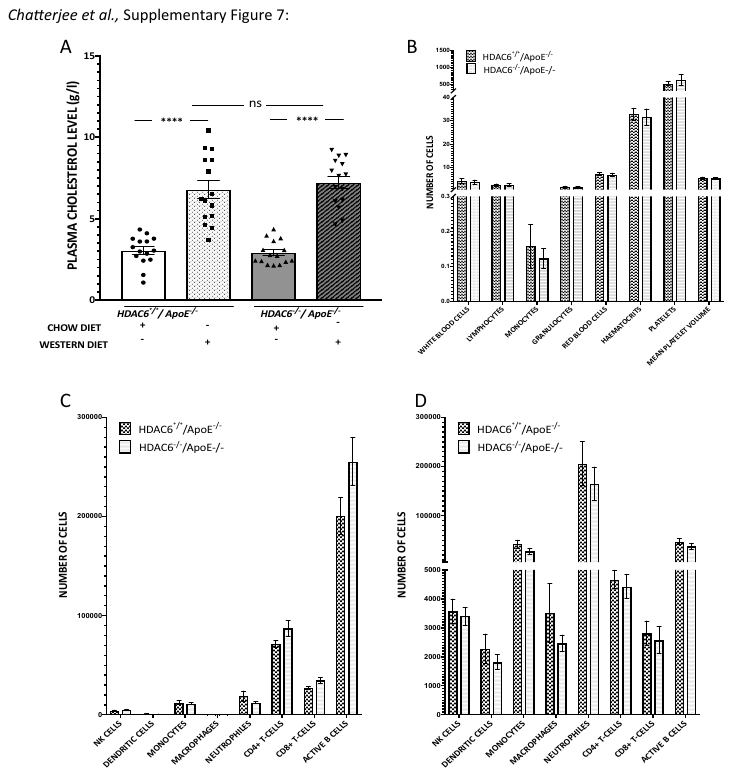
**Supplementary figure 8: Deletion of *HDAC6* *in vivo* does not affect plasma cholesterol levels and immune cell profile.** Chimeric *HDAC6^-/-^/ApoE^-/-^* mice and littermate controls *HDAC6^+/+^/ApoE^-/-^* transplanted with *HDAC6^+/+^/ApoE^-/-^* bone marrow were fed with a high fat diet for 10 weeks. **(A)** Analysis of plasma cholesterol levels compared between the two groups, before and after the Western diet (*HDAC6^+/+^/ ApoE-/-* , n=15; *HDAC6-/- ApoE-/-,* n=15). **(B)** Number of cells for different populations in the blood of *HDAC6^+/+^/ ApoE-/-* (n=4) and *HDAC6-/- ApoE-/-* (n=5) mice after 10 weeks of high fat diet. Comparison of inflammatory cell populations in the spleen **(C)** and bone marrow **(D)** of *HDAC6^+/+^/ ApoE-/-* (n=6) and *HDAC6-/- ApoE-/- (*n=6). Data represent means ± SEM (Mann-Whitney test).
